# Negative controls of chemical probes can be misleading

**DOI:** 10.1101/2020.09.30.320465

**Authors:** Jinyoung Lee, Matthieu Schapira

## Abstract

Chemical probes are selective modulators that are used in cell assays to link a phenotype to a gene and have become indispensable tools to explore gene function and discover therapeutic targets. While binding to off-targets can be acceptable or beneficial for drugs, it is a confounding factor for chemical probes, as the observed phenotype may be driven by inhibition of an unknown off-target instead of the targeted protein. A negative control – a close chemical analog of the chemical probe that is inactive against the intended target – is typically used to verify that the phenotype is indeed driven by targeted protein. Here, we compare the selectivity profiles of four unrelated chemical probes and their respective negative controls and find that the control is sometimes inactive against up to 80% of known off-targets, suggesting that a lost phenotype upon treatment with the negative control may be driven by loss of inhibition of the off-target. To extend this analysis, we inspect the crystal structures of 90 pairs of unrelated proteins, where both proteins within each pair is in complex with the same drug-like ligand, and estimate that in 50% of cases, methylation (a simple chemical modification often used to generate negative controls) of the ligand at a position that will preclude binding to one protein (intended target) will also preclude binding to the other (off-target). These results uncover a risk associated with the use of negative controls to confirm gene-phenotype associations. We propose that a best practice should rather be to verify that two chemically unrelated chemical probes targeting the same protein lead to the same phenotype.

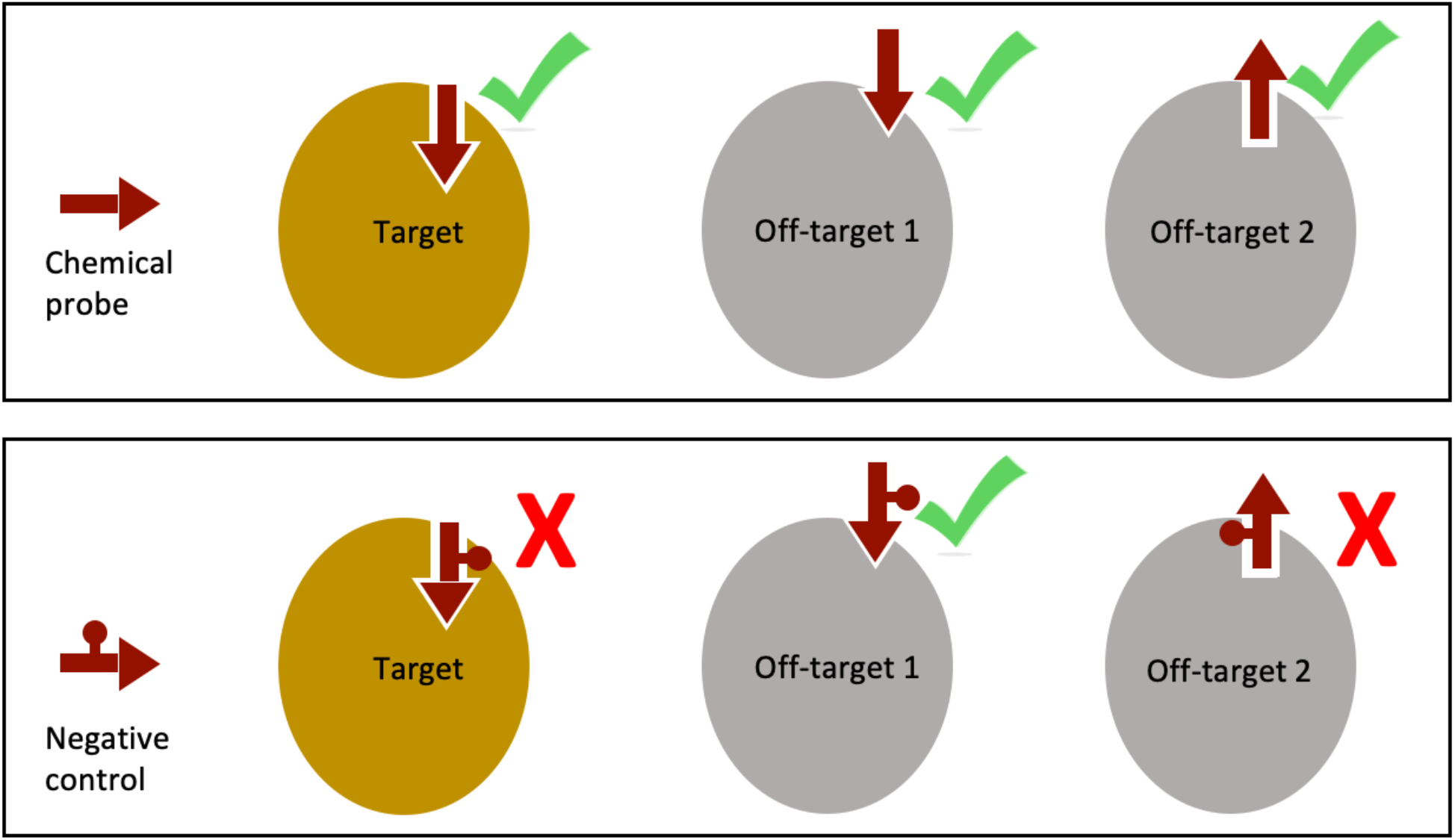

---

Chemical probes are small molecules ligands that are used to modulate the function of a specific biomolecular target – generally a protein – in cells. Along with antibodies, they are among the most powerful tools to explore gene function and disease association and are instrumental in modern day chemical biology and drug discovery^1–3^. Because chemical probes are used to link a phenotype to a gene (if a phenotype is observed upon treatment with the chemical probe, it is attributed to the protein targeted by the probe) specificity is an essential attribute of chemical probes. Compounds of poor quality or used at inappropriate dosage can inhibit other proteins than the intended target and resources such as chemicalprobes.org, Probe Miner or Probes & Drugs are developed to assist in the proper selection and use of chemical probes ^4–6^.

To limit the risk of wrongly attributing a phenotype to the targeted protein while it may be driven by inhibition of an unknown off-target, a recommended practice is to use a negative control. These compounds are close chemical analogs of the probe that are designed to be inactive against the protein targeted by the chemical probe, but expected to remain active against off-targets: if a phenotype is observed upon treatment with a chemical probe, but absent when using its negative control, the phenotype is confidently associated to the protein for which the chemical probe was designed^4^. A critical but unevaluated assumption here is that the chemical modification engineered into the negative control to abrogate binding to the targeted protein has no effect on off-targets: if the negative control no longer binds to an off-target, a phenotype will be present upon treatment with the chemical probe, absent upon treatment with its negative control, and may be driven by inhibition of the unknown off-target while attributed to inhibition of the targeted protein, leading to the wrong gene-phenotype association.

To evaluate the risk associated with the use of negative controls, we first compare the selectivity profile of four well characterized chemical probes with that of their respective negative controls and observed a significant loss of activity of negative controls against off-targets. To expand the analysis to more compounds, we extracted multiple pairs of unrelated proteins bound to identical small-molecule ligands from the Protein Data Bank (PDB) and calculated the risk that a chemical modification affecting binding of the ligand to one protein will affect binding to the other protein. Our results support the notion that the use of negative controls cannot always reliably confirm gene-phenotype association. We rather encourage an alternative practice, which is to replicate a phenotype using two chemical probes with distinct scaffolds but targeting the same protein.

To evaluate whether negative controls retain binding to off-targets of chemical probes, we compared the experimental profiling data of four chemical probes and their respective negative controls (**Table 1**, **Supplementary Table 1**). The chemical probes are A-196, targeting the methyltransferases SUV420H1 and SUV420H2^7^, NVS-MLLT-1, which binds the YEATS domain of MLLT1 and MLLT3^8^, NVS-BPTF-1, a ligand targeting the bromodomain of BPTF^9^, and LLY-507, an inhibitor of the methyltransferase SMYD2^10^. Each chemical probe and its negative control were profiled against an identical panel of GPCRs and other receptors, transporters, kinases and other enzymes (**Supplementary Table 1**)^7–9,11^. We find that in some cases, up to 83% of off-targets inhibited by the chemical probe are not inhibited (or inhibited over 100 fold less potently) by the negative control (**Table 1**). In some cases, the ligands are potent inhibitors of the off-target. For example, NVS-MLLT-1 has an IC_50_ of 250 nM for acetylcholinesterase (ACES) and 290 nM for Histamine receptor H3 but the negative control, which differs by only one heavy atom, is inactive against these proteins. The recommended dosage for NVS-MLLT-1 in cell assays is between 1 and 10 μM^8^. At these concentrations, it may have a significant effect on ACES and H3 which cannot be deconvoluted by the negative control. LLY-507 is a chemical probe that hits numerous off-targets, including M1-M4, NET and SERT, all with Ki values below 800 nM, but the negative control SGC705, which again differs by one heavy atom only, binds to none of these receptors^11^. If treatment with the chemical probe produces a phenotype driven by inhibition of one of these off-targets, the absence of phenotype upon treatment with the negative control may be wrongly attributed to inhibition of the intended target. We note that loss of binding to off-targets by negative controls is unrelated to the quality of the probe or the negative control. For instance, NVS-MLLT-1 is a high-quality chemical probe with very few off-targets and its negative control is almost identical chemically but no longer binds five out of six off-targets. Conversely, the entire 2-cyclopropyl-pyrazole group of NVS-BPTF-1 is shifted by one chemical bond in the negative control of NVS-BPTF-C, but binding is preserved to the four known off-targets (**Table 1**). These experimental profiling data suggest that negative controls of chemical probes may not always be relied upon to validate a putative gene-phenotype association revealed by a chemical probe.

**Table 1:**
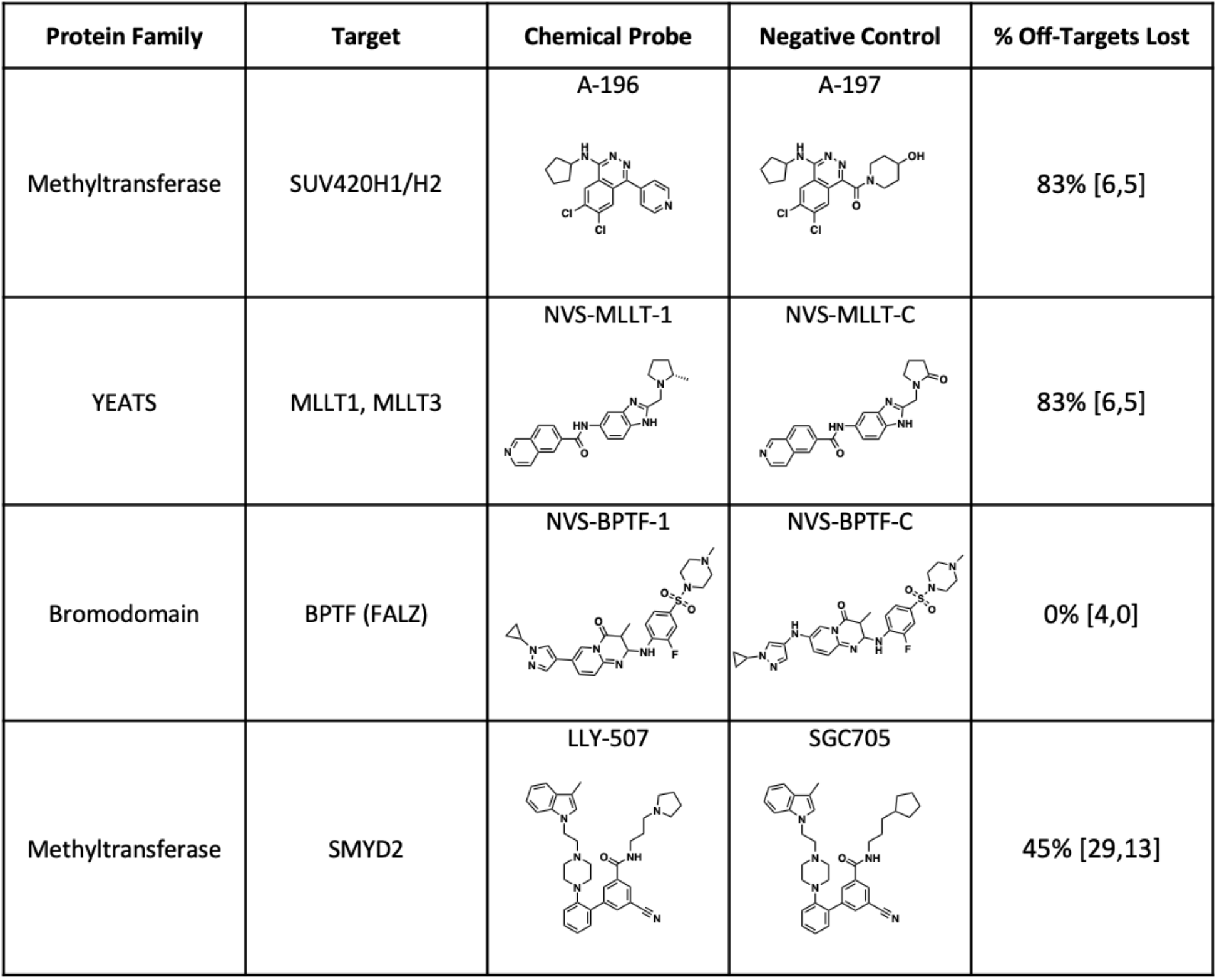
Experimental profiling of chemical probes and their negative control. For each chemical probe, the percent of off-targets that are not inhibited by the negative control is indicated. The number of known off-targets of the chemical probe and the number of off-targets no longer bound by the negative control are shown in brackets. All off-targets are listed in Supplementary Table 1.

| Protein Family | Target | Chemical Probe | Negative Control | % Off-Targets Lost |
| --- | --- | --- | --- | --- |
| Methyltransferase | SUV420H1/H2 | A-196 | A-197 | 83% [6,5] |
| YEATS | MLLT1, MLLT3 | NVS-MLLT-1 | NVS-MLLT-C | 83% [6,5] |
| Bromodomain | BPTF (FALZ) | NVS-BPTF-1 | NVS-BPTF-C | 0% [4,0] |
| Methyltransferase | SMYD2 | LLY-507 | SGC705 | 45% [29,13] |

To extend this analysis to a larger number of compounds and target / off-target pairs, we estimated the risk that a negative control generated for a specific target will be inactive against off-targets using structures available in the PDB. We systematically extracted from the PDB the structures of identical ligands in complex with unrelated protein domains and evaluated whether addition of a methyl group on the compound (a small chemical modification that is often used to generate negative controls) would differently affect binding to each of the targeted proteins. We first verified that the effect of adding a methyl group to a co-crystallized ligand could rapidly and accurately be predicted computationally: methylating the cyclohexyl group of the JAK3 kinase inhibitor FM-381 in ortho position (compound 5) is experimentally well tolerated^12^. Conversely, methylating the pyrrole nitrogen produced the negative control FM-479 that had no significant activity^13^ (**Supplementary Table 2**). We calculated Van der Walls energy associated with the bound ligands before and after methylation (see the methods section for details) and find indeed steric clashes with FM-479 but not with compound 5 or FM-381. Similarly, we determine computationally that methylating the pyrrole nitrogen of BAY-707, an inhibitor of the Nudix protein NUDT1, is not tolerated while adding a methyl group on the methylene bridge of compound 12, an EP300 acetyltransferase inhibitor, has little effect on potency, both of which are confirmed experimentally (**Supplementary Table 2)**^14,15^.

Having verified that the introduction of inactivating steric clashes associated with the methylation of a co-crystallized small-molecule ligand can be determined computationally, we conducted virtual methyl scans on all compounds bound to at least two unrelated proteins in the PDB: each hydrogen atom was sequentially replaced with a methyl group and steric clashes (between the methylated compound and the protein or internal to the bound ligand) associated with the methylation event were evaluated. For instance, the dual JAK2-BRD4 inhibitor TG101209 was crystallized in complex with the kinase domain of JAK2 and the bromodomain of BRD4 (**Figure 1a**). We sequentially replaced each hydrogen of the compound with a methyl group and calculated the resulting energetic penalty due to steric clashes, assuming that the bound conformation remains unchanged after methylation. Out of 25 positions, methylation of five and four on TG101209 are detrimental for binding to JAK2 and BRD4 respectively (**Figure 1b**). Of these sterically constrained hydrogens found in the inhibitor, three are shared between the JAK2-bound and the BRD4-bound states: if one were to make a BRD4 negative control by methylating a position in TG101209 that is sterically constrained in the BRD4-bound state, there would therefore be 75% chances that this position is also sterically constrained in the JAK2-bound state and that the BRD4 negative control would no longer bind to JAK2. In other word, if TG101209 was used as a BRD4 chemical probe and JAK2 was an off target, there is a 75% risk that the negative control is also inactive against the off-target. Inversely, if TG101209 were used as a chemical probe against JAK2, the percent risk (pcR) that a negative control would no longer bind to the off-target (BRD4) would be 60% (**Figure 1b**).

**Figure 1:**
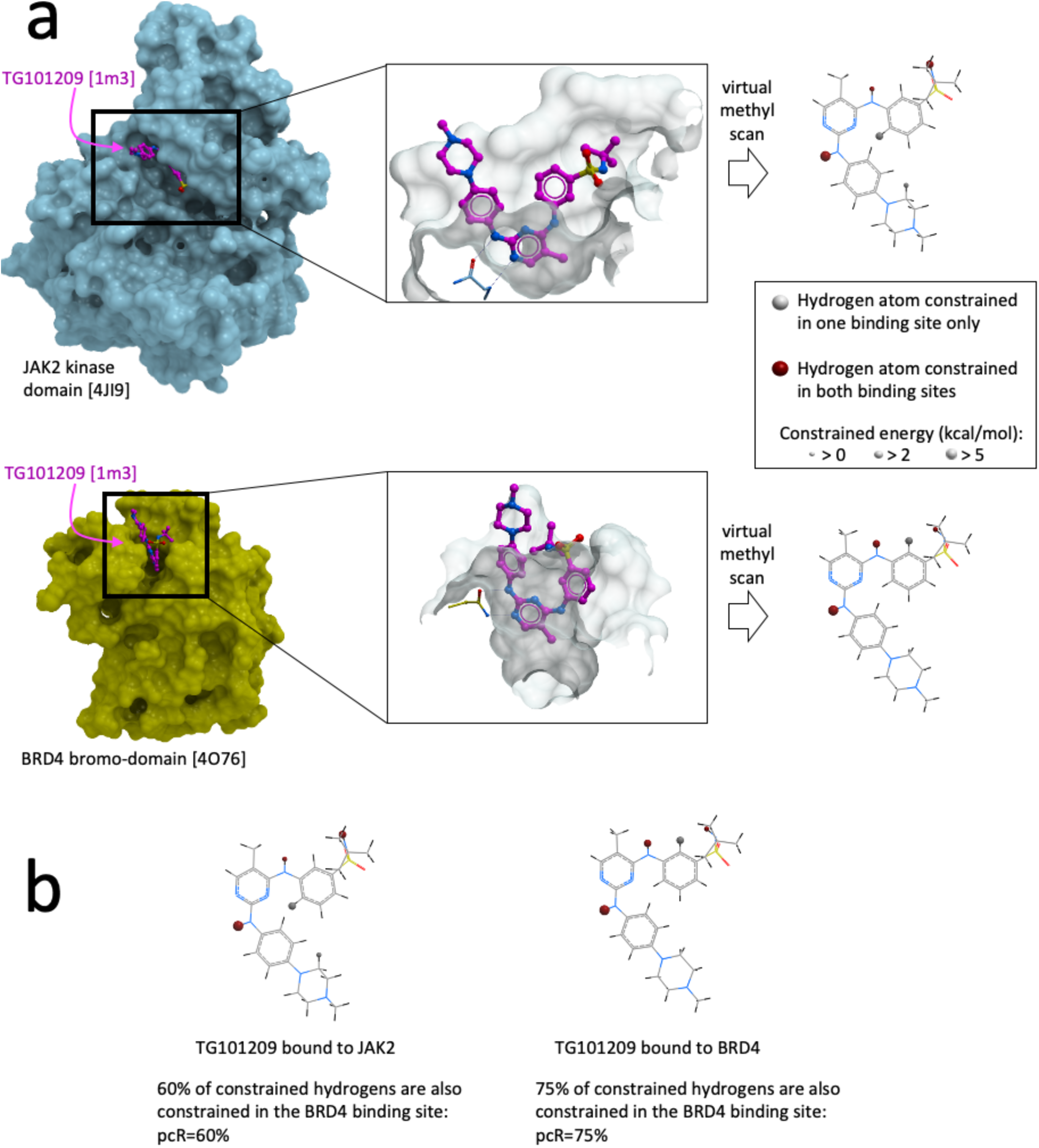
Calculating the risk that a negative control does not bind an off-target. a) Van der Waals energy of the bound ligand associated with the substitution of each hydrogen atom with a methyl group in compound TG101209 (PDB ligand ID 1m3) are calculated using the crystal structures of the compound bound to JAK2 (PDB code 4JI9) and bound to BRD4 (PDB code 4O76). b) Decorating TG101209 with a methyl group to abrogate binding to JAK2 has 60% chances of abrogating binding to BRD4 (percent risk pcR = 60%). Decorating TG101209 with a methyl group to abrogate binding to BRD4 has 75% chances of abrogating binding to JAK2 (pcR = 75%). Because of their structural equivalence, the three hydrogen atoms of a methyl group in the parent compound count as one.

We next extended this analysis to the entire PDB. We found 41 compounds, each bound to at least two unrelated protein domains in the PDB, representing 90 pairs of unrelated proteins (**Supplementary Table 3**). The compounds could be grouped into 17 clusters with structurally distinct chemical scaffolds (**Supplementary Table 4**). For each pair of proteins A and B, we calculated the two pcR values that a negative control for target A would be inactive on target B and that a negative control for target B would be inactive on target A. We find a wide distribution of pcR values across all protein pairs and within each cluster, but risks were consistently large, with an average of 51% and median of 50% (**Figure 2, Supplementary Table 3**).

**Figure 2:**
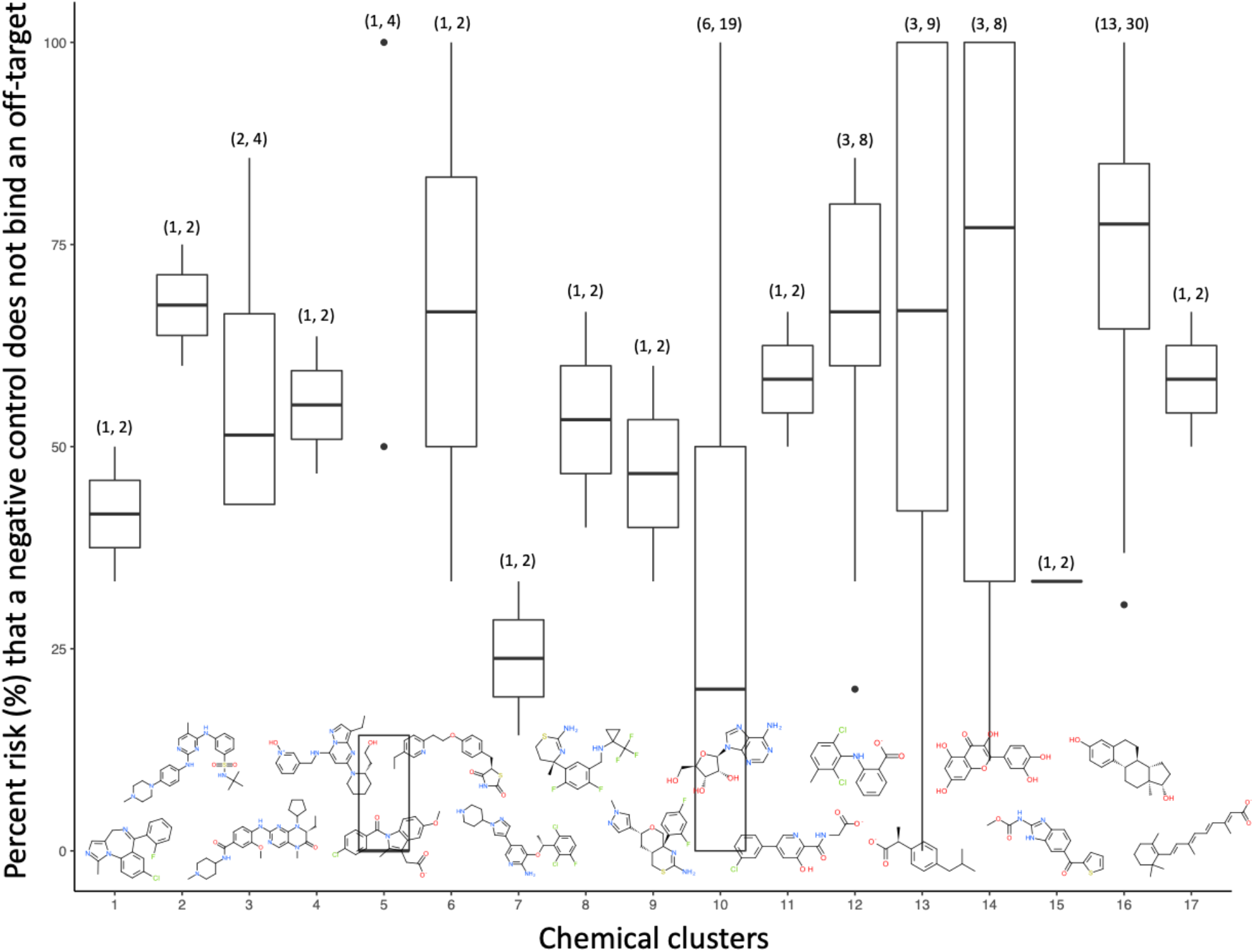
pcR values across 17 chemical clusters representing 41 compounds and 90 pairs of unrelated proteins. Each boxplot represents the distribution of risk factors for all compounds and for all pairs of unrelated proteins for a given chemical cluster. A representative molecule is shown at the bottom; the number of compounds and proteins for each cluster are shown at the top. Additional data and structures are provided in Supplementary Tables 3 and 4.

To make sure that high pcR values would not be driven by structural similarities between protein domains, we only used protein pairs that are structurally as dissimilar as the bromodomain of BRD4 and the kinase domain of JAK2 (protein backbone root mean square deviation [RMSD] within 10 Å of the bound ligand greater than 9.5 Å). We also verified that there is no correlation between RMSD and pcR values (**Supplementary Figure 1a**).

Another possibility is that pcR values would be especially high when ligands are deeply buried into a binding pocket, in which case most hydrogen atoms would be sterically constrained and any methylation necessary to abrogate binding to one protein would abrogate binding to the other. However, we do not see a strong correlation between the percent of hydrogens sterically constrained and pcR values (**Supplementary Figure 1b, Supplementary Table 3**). This is reflected by the dual JAK2-BRD4 inhibitor (**Figure 1**). Only 20% of hydrogens are sterically constrained in JAK2 and 16% in BRD4, but the majority of these constrained hydrogens are the same for the two proteins. There is a strong rationale why the central amino pyrimidine ring of TG101209 engages both JAK2 and BRD4: this pharmacophoric center includes juxtaposed hydrogen-bond donating and accepting groups which can form a canonical dual hydrogen-bond with the hinge region linking the two lobes of JAK2 and with the asparagine at the bottom of the BRD4 binding pocket, conserved across most bromodomains^16,17^. While the two binding pockets are structurally unrelated, they exploit the same interaction hotspot in the ligand, which may be why we found three distinct bromodomain-kinase pairs (**Supplementary Tables 3 and 5**). We find that constrained hydrogens are more evenly distributed across the bound ligand in other protein pairs (**Table 2**, **Supplementary Table 4**), suggesting that the shared interaction hotspot observed between kinases and bromodomains may not be a general rule for poly-pharmacology.

It has been argued that the median number of protein targets per drugs is about 40^18^. Similarly, chemical probes are not expected to be absolutely selective at the proteome scale. While mediocre selectivity may be acceptable or even necessary for a drug, it can be a confounding factor when interpreting phenotypic data produced by a chemical probe. To limit this liability, we and others have encouraged the use of negative controls of chemical probes to confirm that observed phenotypes are driven by inhibition of the intended target. Here, using orthogonal approaches based on experimental profiling data and on structural data, we find that chemical modifications introduced in negative controls have on average 50% chance to affect binding not only to the intended target but also to unknown off-targets.

An alternative approach that is encouraged to validate gene-phenotype association is to verify that the phenotype is reproduced when using two chemically unrelated probes. Indeed, it is extremally unlikely that two compounds with distinct chemical scaffolds would both bind not only the same target but also the same off-target. Developing two unrelated chemical probes for the same target is demanding from a medicinal chemistry perspective, but our data indicates that this additional effort is justified. When available, chemical biologists should consider progressing multiple hits from a primary screening campaign towards the development of more than one chemical probes.

## Online content

Methods, codes and extended data are available as supplementary online information.

## Acknowledgment

The SGC is a registered charity (no. 1097737) that receives funds from AbbVie, Bayer Pharma AG, Boehringer Ingelheim, Canada Foundation for Innovation, Eshelman Institute for Innovation, Genome Canada through Ontario Genomics Institute, Innovative Medicines Initiative (EU/EFPIA) [ULTRA-DD grant no. 115766], Janssen, Merck & Co., Novartis Pharma AG, Ontario Ministry of Economic Development and Innovation, Pfizer, São Paulo Research Foundation-FAPESP, Takeda, and the Wellcome Trust. Additional funding from the Collaborative Research and Training Experience grant (Aled Edwards 432008-2013) from the Natural Sciences and Engineering Research Council of Canada. MS gratefully acknowledges funding from NSERC (grants RGPIN-2019-04416 and ALLRP 555329-20).

