## Supplementary information for "Negative controls of chemical probes can be misleading"

### METHODS

All computational work was conducted with ICM-pro 3.8-7d (Molsoft, San Diego). All codes are provided at the end of this document.

#### Assembling a set of matched protein pairs

To assemble a valid collection of pairs of protein structures in complex with identical ligands, the following conditions were used: (1) co-crystalized ligands must meet Lipinski rule of five; (2) co-crystalized ligands must be occupying a well-defined pocket rather than laying on a flat surface: ligand desolvation value > 70%. These two conditions were addressed as described in our previous work (PMID 32084498). (3) Structures must have a resolution of 3Å or better. (4) Finally, ligands must be co-crystalized with at least two unrelated proteins. To meet this condition, we first kept the structure with the highest resolution when multiple structures of the same protein (defined by its InterPro ID) and the same ligand (defined by its PDB ID) were available. Next, for each pair of protein in complex with the same ligand, we used ICM to calculate backbone RMSD between protein domains occupied by the ligand (defined by aminoacids within 10 Å of the bound ligand). Protein pairs with RMSD values > 9.5Å (based on the RMSD between BRD4 and JAK2 ligand binding domains of 9.8Å) were retained.

#### Energy calculation

The energy of the co-crystalized ligand and the energies of the methylated analogues were calculated for each structure. All calculations were conducted with ICM (Molsoft, San Diego). First, the co-crystallized ligand was tethered to its original binding pose (tzWeight =1) and minimized locally in the internal coordinates space with default energy terms (including Van der Waals (VdW), hydrogen bonds, electrostatics and torsion terms). The Van de Waals energy of the bound ligand was then calculated with the command “show ey” followed by “Energy(“vw,14”)” using a soft Van der Waals energy term (vwMethod = 3). The overall energy calculation process was repeated three times for each structure, and the averages of those three rounds were used for the analysis. Standard deviations were low and are reported in Supp Table 3. If the Van der Waals energy of the un-methylated, energy-minimized ligand was positive, the structure was rejected.

Next, the Van der Waals energy of methylated analogues were calculated as follows. The co-crystallized ligand in its energy-minimized pose was mono-methylated *in silico* with ICM and tethered to the original, unmodified ligand with a high tether weight (tzWeight = 200.) to constrain the conformation of the virtual negative control to the binding pose of the un-methylated crystallized ligand (previously relaxed as described above). The energy of the tethered virtual negative control was globally minimized using ICM's Monte Carlo method (command: montecarlo), followed by a local minimization (command: minimize). The Van der Waals energy of the methylated compound was then calculated as described above. This process of generating and calculating energy of a virtual negative control was repeated for each hydrogen atom of the parent compound. A hydrogen atom was judged as sterically restrained if replacing it with a methyl group resulted in positive Van der Waals energy of the bound ligand.

| Protein Family | Specific Target | Probe | Negative Control (NC) | Probe Off-Target | NC Off-Target | % Off-Target Lost |
|---|---|---|---|---|---|---|
| Methyltransferase | SUV420 H1/H2 | A-196 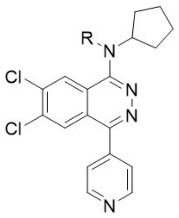 | A-197 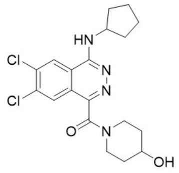 | A1 [Ki 0.021 μM] | A1 [ Ki 3.1 μM] | 83 |
| | | | | A2A [Ki 0.028 μM] | A2A [Ki 20 μM] | |
| | | | | A3 [Ki 2 μM] | A3 [Ki 13 μM] | |
| | | | | Cl- channel (GABA-gated) [Ki 1.8 μM] | n/a | |
| | | | | NK2 [Ki 4.1 μM] | n/a | |
| | | | | delta 2 (DOP) [Ki 9 μM] | n/a | |
| YEATS | MMT1, MLLT3 | NVS-MLLT-1 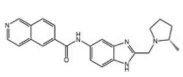 | NVS-MLLT-C 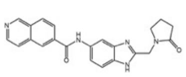 | PIM1 [$IC_{50}$ 8.6 μM] | n/a | 83 |
| | | | | TRB2 [$IC_{50}$ 3.5 μM] | TRB2 [$IC_{50}$ 3.5 μM] | |
| | | | | ACES [$IC_{50}$ 0.25 μM] | n/a | |
| | | | | H3 [$IC_{50}$ 0.29 μM] | n/a | |
| | | | | M2 [$IC_{50}$ 1.8 μM] | n/a | |
| | | | | NET [$IC_{50}$ 8.7 μM] | n/a | |
| Bromodomains | BPTF (FALZ) | NVS-BPTF-1 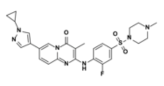 | NVS-BPTF-C 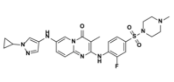 | Ad3 [$IC_{50}$ 4.9 μM] | Ad3 [$IC_{50}$ 13 μM] | 0 |
| | | | | D3 [$IC_{50}$ 3.5 μM] | D3 [$IC_{50}$ 0.95 μM] | |
| | | | | H3 [$IC_{50}$ 4.0 μM] | H3 [$IC_{50}$ 23 μM] | |
| | | | | ACES [$IC_{50}$ 30 μM] | ACES [$IC_{50}$ 5.5 μM] | |
| | | | | n/a | COX2[$IC_{50}$ 0.8 μM] | |
| Methyltransferase | SMYD2 | LLY-507 (Multiple Off-target Effects) 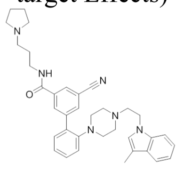 | SGC705 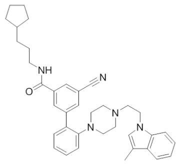 | 5-HT1A [Ki 0.12 μM] | 5-HT1A [$IC_{50}$ 0.14 μM] | 45 |
| | | | | 5-HT1B [Ki 1.2 μM] | n/a | |
| | | | | n/a | 5-HT2A [$IC_{50}$ 0.07 μM] | |
| | | | | 5-HT2B [Ki 2.2 μM] | 5-HT2B [$IC_{50}$ 1.4 μM] | |
| | | | | 5-HT2C [Ki 1.9 μM] | 5-HT2C [$IC_{50}$ 1.5 μM] | |
| | | | | 5-HT5a [Ki 1.9 μM] | n/a | |
| | | | | 5-HT6 [Ki 1.4 μM] | 5-HT6 [Ki 2.2 μM] | |
| | | | | 5-HT7 [Ki 0.3 μM] | 5-HT7 [Ki 2.5 μM] | |
| | | | | Alpha1A [Ki 0.24 μM] | Alpha1A [Ki 0.82 μM] | |
| | | | | Alpha1B [Ki 0.91 μM] | Alpha1B [Ki 3.5 μM] | |
| | | | | Alpha2A [Ki 0.66 μM] | Alpha2A [Ki 0.52 μM] | |

| | | | | | |
|---|---|---|---|---|---|
| | | | | Alpha2B [Ki 1.3 µM] | Alpha2B [Ki 0.14 µM] |
| | | | | Alpha2C [Ki 0.029 µM] | Alpha2C [Ki 0.32 µM] |
| | | | | D2 [Ki 2.5 µM] | n/a |
| | | | | D3 [Ki 2.9 µM] | n/a |
| | | | | D4 [Ki 2.9 µM] | n/a |
| | | | | n/a | D5 [Ki 3.2 µM] |
| | | | | H3 [Ki 0.69 µM] | n/a |
| | | | | KOR [Ki 0.022 µM] | KOR [$IC_{50}$ 2.3 µM] |
| | | | | M1 [Ki 0.57 µM] | n/a |
| | | | | M2 [Ki 0.74 µM] | n/a |
| | | | | M3 [Ki 0.59 µM] | n/a |
| | | | | M4 [Ki 0.45 µM] | n/a |
| | | | | MOR [Ki 0.35 µM] | MOR [$IC_{50}$ 0.58 µM] |
| | | | | NET [Ki 0.57 µM] | n/a |
| | | | | SERT [Ki 0.5 µM] | n/a |
| | | | | Sigma 1 [Ki 0.13 µM] | Sigma 1 [Ki 1.7 µM] |
| | | | | Sigma 2 [Ki 0.059 µM] | Sigma 2 [Ki 0.46 µM] |
| | | | | n/a | PBR [Ki 0.69 µM] |

**Supplementary Table 1: Profiling data for four chemical probes and their negative controls**. Only proteins where compounds are active are shown. NVS-BPTF-1 and its negative control NVS-BPTF-C were profiled on a panel from the Novartis Institute of Biomedical Research consisting of 15 GPCRs, 3 transporters, 3 nuclear hormone receptors, 48 kinases and 7 enzymes[1]. NVS-MLLT-1 and its negative control NVS-MLLT-C were profiled against 59 kinases, 7 additional enzymes, 16 GPCRs and 5 other diverse proteins[2]. A-196 and A-197 were profiled against 54 GPCRS and transporters from CEREP[3]. LLY-507 and SGC-705 were profiled against of 45 GPCRs and transporters from the NIMH Psychoactive Drug Screening Program[4].

| Chemical probe | | | Methylated analogue | | | Methylated analog is active | Methylated analog is predicted active |
|---|---|---|---|---|---|---|---|
| Structure | $IC_{50}$ (nM) | VdW energy (kcal/mol) | Structure | $IC_{50}$ (nM) | VdW energy (kcal/mol) | | |
| **FM-381** Compound 4 (PDB: 5lwm) 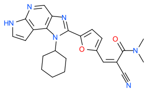 | 0.127 | -8.327 ± 0. | **Compound 5** (PDB: 5lwn) 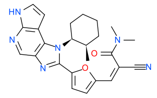 | 0.154 | -5.915 ± 0.02 | yes | yes |
| **FM-381** Compound 4 (PDB: 5lwm) 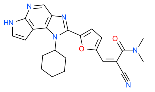 | 0.127 | -8.327 ±0. | **FM-479** 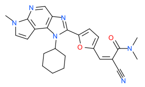 | >1,000 | 0.981 ± 0.00 | no | no |
| **BAY-707** (PDB: 5nhy) 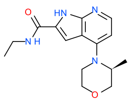 | 2.3 | -9.428 ± 0. | **BAY-604** 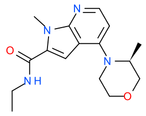 | >20,000 | 9.649 ± 0.03 | no | no |
| **CPI-1612** Compound 17 (PDB: 6v8n) 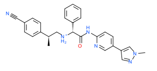 | 8 | -15.39 ± 0. | **Compound 12** (PDB: 6v90) 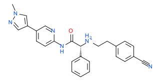 | 19 | -9.585 ± 0.8 | yes | yes |

**Supplementary Table 2**: **Experimentally measured and computationally predicted effects of methylating a chemical probe**. Compound structures, experimental $IC_{50}$ values and computationally calculated Van der Waals clashes for the bound ligand (VdW energy) are shown for the JAK3 kinase ligands FM-381, compound 5 and FM-479[5,6], the NUDT1 ligands BAY-707 and BAY-604[7] and the EP300 ligands CPI-1612 and compounds 12[8].

| **Cluster** ($N_{Ligand}$, $N_{PDB}$); **Example Ligand** | **Example PDB$_1$** | **Example PDB$_2$** |
|---|---|---|
| 1 (1, 2); 08j 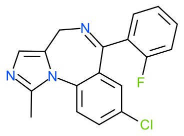 | 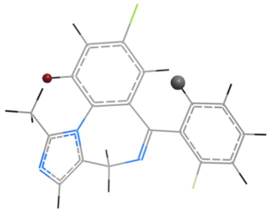 Bromodomain; BRD4; 3u5k; 50% | 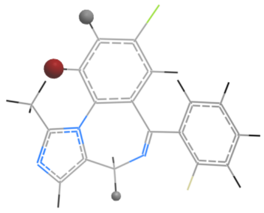 Cytochrome P450; CP3A4; 5te8; 33% |
| 2 (1, 2); 1m3 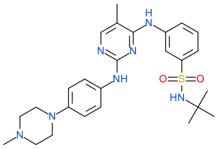 | 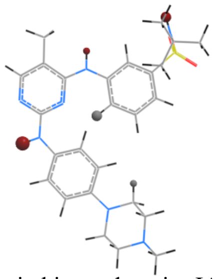 Protein kinase domain; JAK2; 4ji9; 60% | 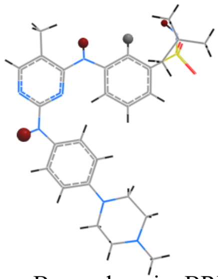 Bromodomain; BRD4; 4o76; 75% |
| 3 (2, 4); r78 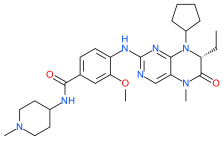 | 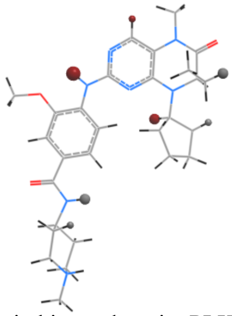 Protein kinase domain; PLK1; 2rku; 43% | 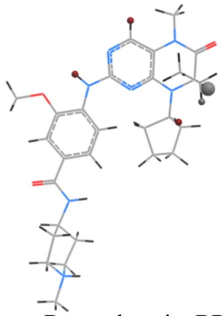 Bromodomain; BRD4; 4ogi; 60% |
| 4 (1, 2); 1qk 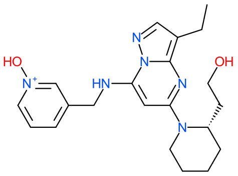 | 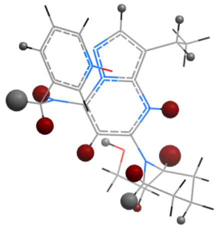 Protein kinase domain; CDK2; 4kd1; 47% | 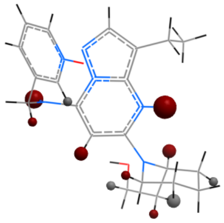 Bromodomain; BRD4; 4o70; 64% |
| 5 (1, 4); imn 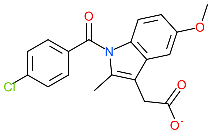 | 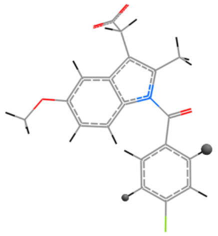 Glutathione S-transferase; PGES2; 1z9h; 0% | 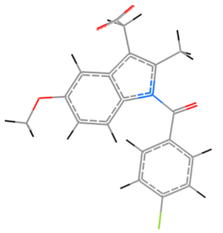 Serum albumin, N-terminal; ALBU; 2bxk; N/A |

| | | |
|---|---|---|
| 6 (1, 2); p1b<br>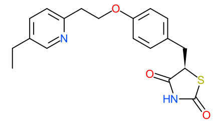 | 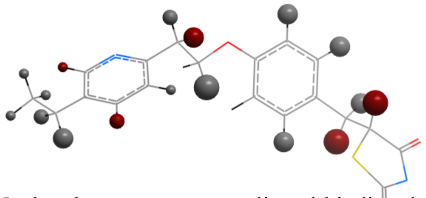<br>Nuclear hormone receptor, ligand-binding domain; PPARG; 2xkw; 33% | 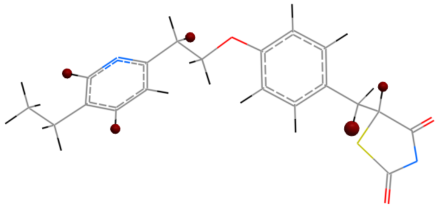<br>Amine oxidase; AOFB; 4a79; 100% |
| 7 (1, 2); vgh<br>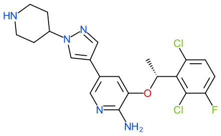 | 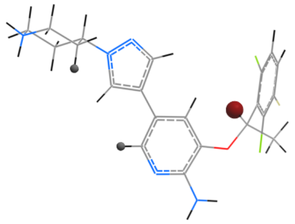<br>Protein kinase domain; ALK; 2yfx; 33% | 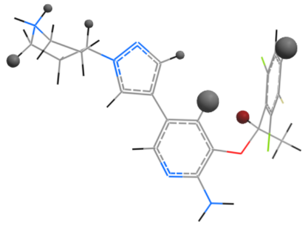<br>NUDIX hydrolase domain; 8ODP; 4c9w; 14% |
| 8 (1, 2); p6u<br>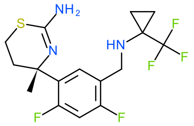 | 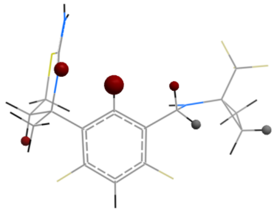<br>Peptidase family A1 domain; BACE1; 5t1u; 67% | 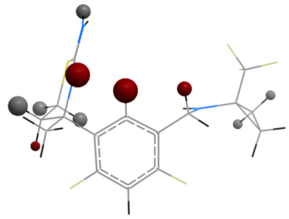<br>Cytochrome P450; CP2D6; 5tft; 40% |
| 9 (1, 2); si5<br>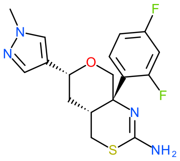 | 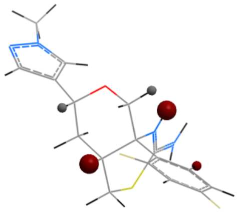<br>Peptidase family A1 domain; BACE1; 4xxs; 60% | 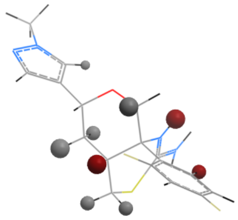<br>Cytochrome P450; CP2D6; 4xry; 33% |
| 10 (6, 19); adn<br>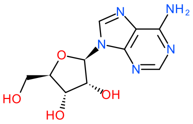 | 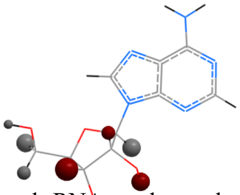<br>Aminoacyl-tRNA synthase, class II (G/P/S/T); SYEP; 4k87; 50% | 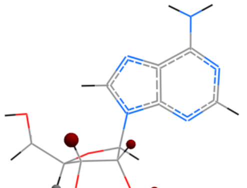<br>S-adenosylmethionine synthase; METK2; 6fcd; 80% |
| 11 (1, 2); a1z<br>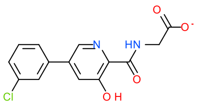 | 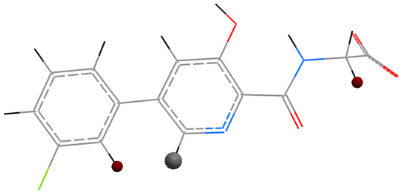<br>Cupin-like domain 8; HIF1N; 5opc; 67% | 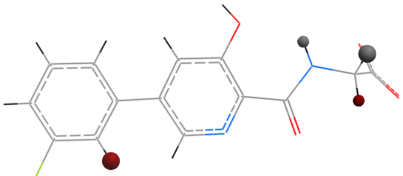<br>Oxoglutarate/iron-dependent dioxygenase; EGLN1; 5ox6; 50% |

| | | |
|---|---|---|
| 12 (3, 8); jms 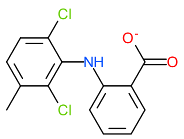 | 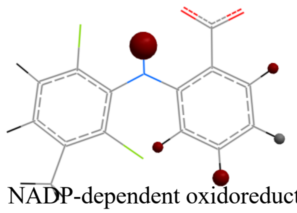 NADP-dependent oxidoreductase domain; AK1C3; 3r6i; 80% | 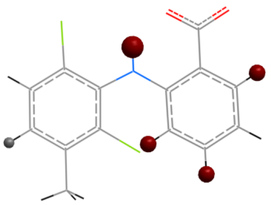 Haem peroxidase, animal type; PGH2; 5ikq; 80% |
| 13 (3, 9); ibp 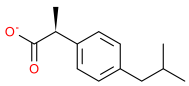 | 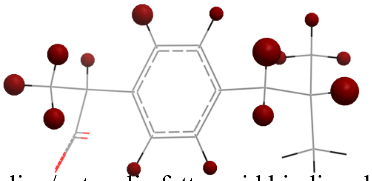 Lipocaline/cytosolic fatty-acid binding domain; FABP4; 3p6h; 100% | 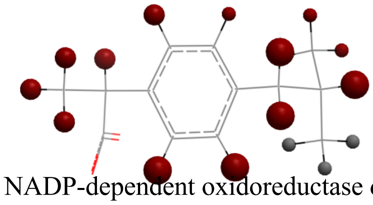 NADP-dependent oxidoreductase domain; AK1C2; 4jtr; 91% |
| 14 (3, 8); que 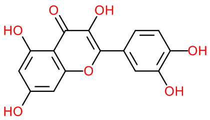 | 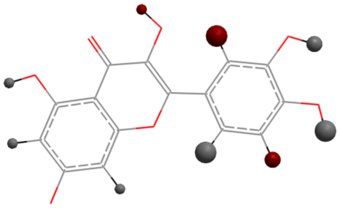 Protein kinase domain; DAPK1; 5auw; 33% | 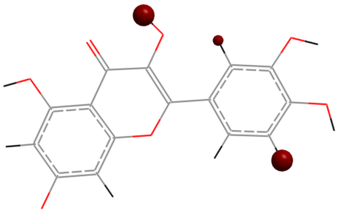 Serine protease, trypsin domain; UROK; 5xg4; 100% |
| 15 (1, 2); nzo 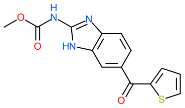 | 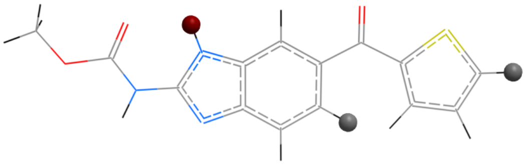 Glutathione S-transferase; HPGDS; 3ee2; 33% | 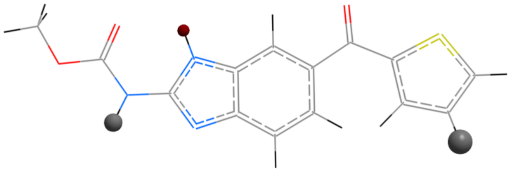 Tubulin/FtsZ, GTPase domain; TBB2B; 5ca1; 33% |
| 16 (13, 30); est 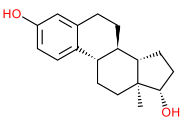 | 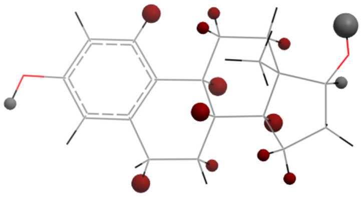 Laminin G domain; SHBG; 1lhu; 79% | 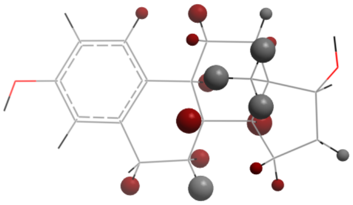 Sulfotransferase domain; ST1E1; 4jvl; 73% |
| 17 (1, 2); rea 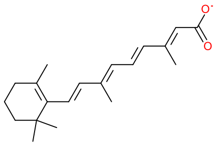 | 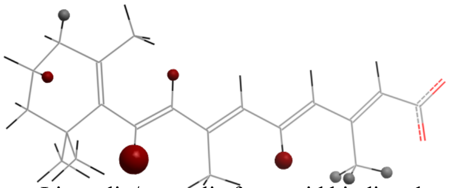 Lipocalin/cytosolic fatty-acid binding domain; RABP2; 2fr3; 67% | 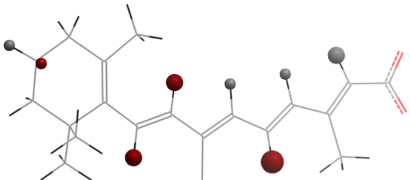 Nuclear hormone receptor, ligand-binding domain; RARG; 2lbd; 50% |

**Supplementary Table 4: pcR across 17 chemical clusters.** Representative compounds are shown for the 17 chemical clusters composed of molecules each found in two or more structures in the PDB in complex with unrelated protein domains. For each cluster, the number of compounds and number of proteins is shown in parenthesis, followed by the PDB ligand name of the representative compound (first column). For each chemical cluster, a representative compound is shown and constrained hydrogens are highlighted as in Figure 2 when bound to two unrelated proteins. The percent risk (pcR) that the negative control of a chemical probe binding to one protein do not bind to the other protein is shown. The full list of 41 compounds found in 90 protein pairs is available in supplementary Table 3.

| **Domain Pair**<br><br>(Domain 1; Domain 2) | **Ligand Cluster** | | | | | | | | | | | | | | | | |
|---|---|---|---|---|---|---|---|---|---|---|---|---|---|---|---|---|---|
| | **1** | **2** | **3** | **4** | **5** | **6** | **7** | **8** | **9** | **10** | **11** | **12** | **13** | **14** | **15** | **16** | **17** |
| NUDIX hydrolase domain; Protein kinase domain | | | | | | | | | | | | ■ | | | | | |
| G protein-coupled receptor, rhodopsin-like; Protein kinase domain | | | | | | | | | | | | | | | ■ | | |
| G protein-coupled receptor, rhodopsin-like; Histidine triad (HIT) protein | | | | | | | | | | | | | | | ■ | | |
| G protein-coupled receptor, rhodopsin-like; Aminoacyl-tRNA synthetase, class II (G/ P/ S/T) | | | | | | | | | | | | | | | ■ | | |
| G protein-coupled receptor, rhodopsin-like; Histidine kinase/HSP90-like ATPase | | | | | | | | | | | | | | | ■ | | |
| G protein-coupled receptor, rhodopsin-like; S-adenosyl-L-homocysteine hydrolase, NAD binding domain | | | | | | | | | | | | | | | ■ | | |
| G protein-coupled receptor, rhodopsin-like; S-adenosylmethionine synthetase, central domain | | | | | | | | | | | | | | | ■ | | |
| G protein-coupled receptor, rhodopsin-like; Histone-lysine N-methyltransferase DOT1 domain | | | | | | | | | | | | | | | ■ | | |
| Nuclear hormone receptor, ligand-binding domain; Lipocalin/cytosolic fatty-acid binding domain | | | | | | | | | | | | | | | | | ■ |
| Nuclear hormone receptor, ligand-binding domain; Oxysterol-binding protein | | | | | | | | | | | | | | | | ■ | |
| Nuclear hormone receptor, ligand-binding domain; Protein kinase domain | | | | | ■ | | | | | | | | | | | | |
| Nuclear hormone receptor, ligand-binding domain; Sulfotransferase domain | | | | | | | | | | | | | | | | ■ | |
| Nuclear hormone receptor, ligand-binding domain; Cytochrome P450 | | | | | | | | | | | | | | | | ■ | |
| Nuclear hormone receptor, ligand-binding domain; Laminin G domain | | | | | | | | | | | | | | | | ■ | |
| Nuclear hormone receptor, ligand-binding domain; Amine oxidase | | | | | | | | | | | ■ | | | | | | |
| Nuclear hormone receptor, ligand-binding domain; MD-2-related lipid-recognition domain | | | | | | | | | | | | | | | | ■ | |
| Nuclear hormone receptor, ligand-binding domain; Glutathione S-transferase | | | ■ | | | | | | | | | | | | | | |
| Nuclear hormone receptor, ligand-binding domain; Serum albumin, N-terminal | | | ■ | | | | | | | | | | | | | | |
| Nuclear hormone receptor, ligand-binding domain; NADP-dependent oxidoreductase domain | | | ■ | | | | | | | | | | | | | ■ | |
| Nuclear hormone receptor, ligand-binding domain; Serpin domain | | | | | | | | | | | | | | | | ■ | |
| Nuclear hormone receptor, ligand-binding domain; Niemann-Pick C1, N-terminal | | | | | | | | | | | | | | | | ■ | |

| |
|---|
| Lipocalin/cytosolic fatty-acid binding domain; AMP-dependent synthetase/ligase |
| Lipocalin/cytosolic fatty-acid binding domain; NADP-dependent oxidoreductase domain |
| Oxysterol-binding protein; Cytochrome P450 |
| Fibrillarin; Nucleoside phosphorylase domain |
| Protein kinase domain; Glycosyl transferase, family 35 |
| Protein kinase domain; Serine proteases, trypsin domain |
| Protein kinase domain; Histidine triad (HIT) protein |
| Protein kinase domain; Bromodomain |
| Protein kinase domain; Aminoacyl-tRNA synthetase, class II (G/ P/ S/T) |
| Protein kinase domain; Heat shock protein 70 family |
| Protein kinase domain; S-adenosyl-L-homocysteine hydrolase, NAD binding domain |
| Protein kinase domain; S-adenosylmethionine synthetase, central domain |
| Protein kinase domain; Histone-lysine N-methyltransferase DOT1 domain |
| Protein kinase domain; Enoyl-CoA hydratase/isomerase, HIBYL-CoA-H type |
| Glycosyl transferase, family 35; Serine proteases, trypsin domain |
| Glycosyl transferase, family 35; Enoyl-CoA hydratase/isomerase, HIBYL-CoA-H type |
| Nucleoside phosphorylase domain; Deoxynucleoside kinase domain |
| Nucleoside phosphorylase domain; PRMT5 arginine-N-methyltransferase |
| Sulfotransferase domain; Laminin G domain |
| AMP-dependent synthetase/ligase; NADP-dependent oxidoreductase domain |
| Cytochrome P450; Bromodomain |
| Cytochrome P450; MD-2-related lipid-recognition domain |
| Cytochrome P450; Glutathione S-transferase |
| Cytochrome P450; Serum albumin, N-terminal |
| Cytochrome P450; Haem peroxidase, animal type |
| Cytochrome P450; NADP-dependent oxidoreductase domain |
| Cytochrome P450; Peptidase family A1 domain |
| Serine proteases, trypsin domain; Enoyl-CoA hydratase/isomerase, HIBYL-CoA-H type |

| | | | | | | | | | | | | | | | | |
|---|---|---|---|---|---|---|---|---|---|---|---|---|---|---|---|---|
| Histidine triad (HIT) protein; Aminoacyl-tRNA synthetase, class II (G/ P/ S/T) | | | | | | | | | | | | | | ■ | | |
| Histidine triad (HIT) protein; Heat shock protein 70 family | | | | | | | | | | | | | | ■ | | |
| Histidine triad (HIT) protein; S-adenosyl-L-homocysteine hydrolase, NAD binding domain | | | | | | | | | | | | | | ■ | | |
| Adenosine/AMP deaminase domain; Deoxynucleoside kinase domain | | | | | | | | | | | | | | ■ | | |
| Aminoacyl-tRNA synthetase, class II (G/ P/ S/T); Heat shock protein 70 family | | | | | | | | | | | | | | ■ | | |
| Aminoacyl-tRNA synthetase, class II (G/ P/ S/T); S-adenosyl-L-homocysteine hydrolase, NAD binding domain | | | | | | | | | | | | | | ■ | | |
| Aminoacyl-tRNA synthetase, class II (G/ P/ S/T); S-adenosylmethionine synthetase, central domain | | | | | | | | | | | | | | ■ | | |
| Aminoacyl-tRNA synthetase, class II (G/ P/ S/T); Histone-lysine N-methyltransferase DOT1 domain | | | | | | | | | | | | | | ■ | | |
| Tubulin/FtsZ, GTPase domain; Glutathione S-transferase | | | | ■ | | | | | | | | | | | | |
| Glutathione S-transferase; Serum albumin, N-terminal | | | ■ | | | | | | | | | | | | | |
| Glutathione S-transferase; NADP-dependent oxidoreductase domain | | | ■ | | | | | | | | | | | | | ■ |
| Oxoglutarate/iron-dependent dioxygenase; Cupin-like domain 8 | | ■ | | | | | | | | | | | | | | |
| Heat shock protein 70 family; S-adenosyl-L-homocysteine hydrolase, NAD binding domain | | | | | | | | | | | | | | | ■ | |
| Heat shock protein 70 family; S-adenosylmethionine synthetase, central domain | | | | | | | | | | | | | | | ■ | |
| Heat shock protein 70 family; Histone-lysine N-methyltransferase DOT1 domain | | | | | | | | | | | | | | | ■ | |
| Serum albumin, N-terminal; NADP-dependent oxidoreductase domain | | | ■ | | | | | | | | | | | | | |
| S-adenosyl-L-homocysteine hydrolase, NAD binding domain; S-adenosylmethionine synthetase, central domain | | | | | | | | | | | | | | | ■ | |
| S-adenosyl-L-homocysteine hydrolase, NAD binding domain; Histone-lysine N-methyltransferase DOT1 domain | | | | | | | | | | | | | | | ■ | |
| Haem peroxidase, animal type; NADP-dependent oxidoreductase domain | | | | | | ■ | ■ | | | | | | | | | |
| Haem peroxidase, animal type; Alpha-ketoglutarate-dependent dioxygenase FTO, catalytic domain | | | | | | ■ | | | | | | | | | | |
| Haem peroxidase, animal type; YAP binding domain | | | | | | ■ | | | | | | | | | | |
| NADP-dependent oxidoreductase domain; Alpha-ketoglutarate-dependent dioxygenase FTO, catalytic domain | | | | | | ■ | | | | | | | | | | |
| NADP-dependent oxidoreductase domain; YAP binding domain | | | | | | ■ | | | | | | | | | | |

**Supplementary Table 5**: For each pair of unrelated protein domains, chemical clusters where at least one ligand is found in complex with both domains in the PDB are highlighted.

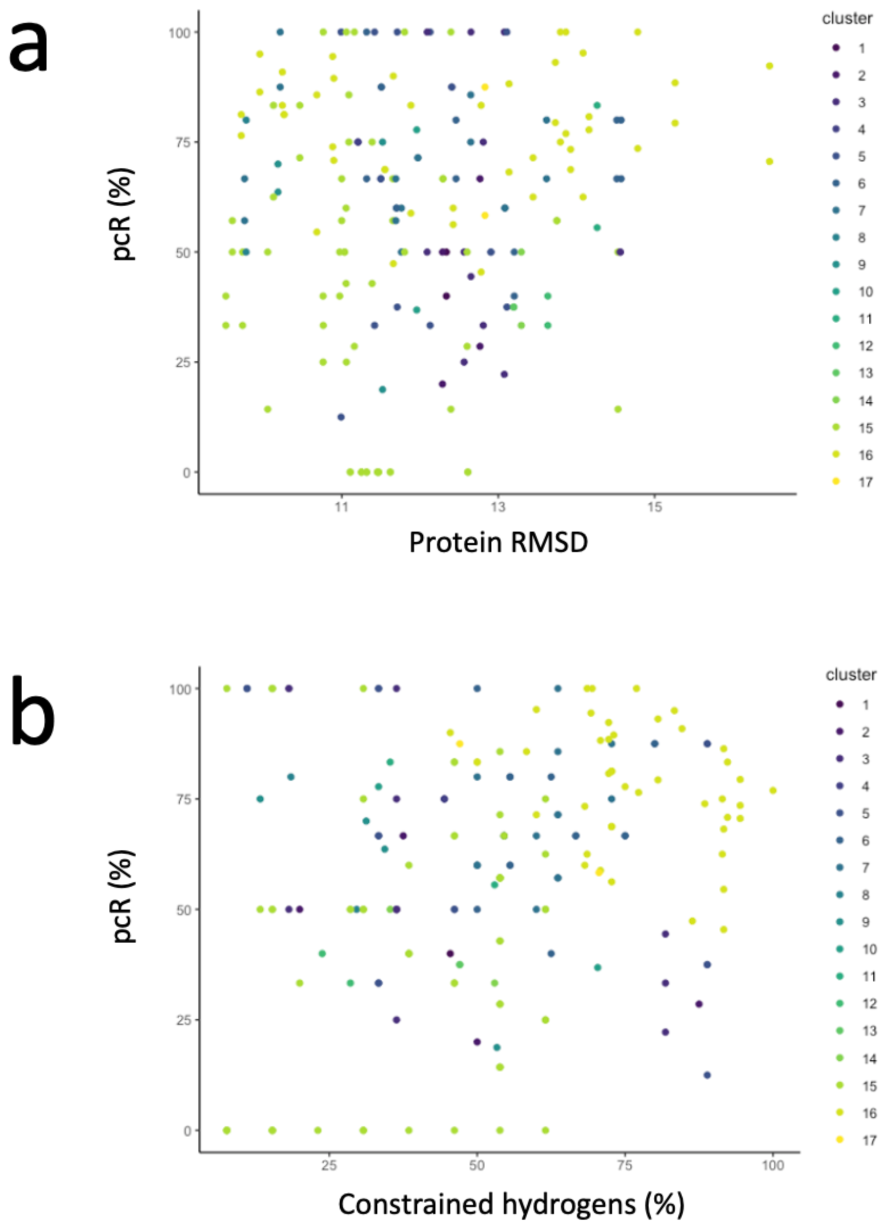

**Supplementary Figure 1**: pcR values do not correlate with the structural similarity between proteins domains (a) or the percentage of hydrogens constrained in a binding site (b).

### Computer Codes

All codes are shown below and can be run with ICM-Pro (Molsoft, San Diego) available from molsoft.com.

<u>1- Data Filter</u>

```
errorAction = 3

# Purpose: To filter the "drug-like" ligand data based on following criteria:
#           1) desolvaiton value
#           2) different domains - identified based on different InterPro IDs
# Input: all_ip_with_pdb (drug_like ligands last updated Dec 17, 2019), pdb_resolu (table)
```

```
# Output: lig_ips

#-------------------------------------------------------------------------------------------------------------
# INPUT VARIABLES
t = all_ip_with_pdb
#-------------------------------------------------------------------------------------------------------------

#-----------------------------------------------------------------------------------------------
function FilterIP( s )
# Prerequisite - this function needs table "pdb_resolu" and table "t" prepared
# Purpose - takes in ligand_name "s" as an argument and returns table "t_final" if and only if the
#           s is co-crystalized with more than one domain (identified using InterPro ID comparisons)

  t_temp = t.lig_name == s
  sort(t_temp.ip_code)
  j = 1
  while j <= Nof(t_temp)
    r_ip = t_temp.ip_code[j]
    t_ip = t_temp.ip_code==r_ip

    # MULTIPLE STRUCTURES FOR ONE DOMAIN - if there are multiple structures within
    #          the same domain, then the structure with the best resolution is retained
    if Nof(t_ip) > 1 then
      add column t_ip Rarray(Nof(t_ip)) name = "resol"
      for jj = 1, Nof(t_ip)
        i_resol = Index(pdb_resolu.pdb_code t_ip.pdb_code[jj])
        t_ip.resol[jj] = pdb_resolu.resolution[i_resol]
      endfor
      delete t_ip.resol == -1            #remove NMR structure
      sort(t_ip.resol)
      lowest_resol = t_temp.pdb_code == t_ip.pdb_code[1]
      if Nof(lowest_resol) != 1 then  #the one with the best resolution
        add t_final lowest_resol[1]
      else
        add t_final lowest_resol
      endif
    else
      add t_final t_ip[1]
    endif
    j = Index(t_temp.ip_code r_ip) + Nof(t_temp.ip_code == r_ip)
    delete t_ip, r_ip
  endwhile
  delete t_temp
  if Nof(t_final) > 1 then               #check if t_final has more than one domains
      return t_final
  endif

endfunction
#-----------------------------------------------------------------------------------------------

# MAIN SCRIPT STARTS:

# STEP 0 - preparing the table
# Note - STEP 0 handles exceptions that are manually implemented for the initial data used in my analysis.
#          If you are using the same initial table as my analysis, keep these exceptions.
#          If not, please check them to ensure they are applicable for your analysis as well.
#          You can add or remove any of the exceptions in STEP 0 based on the data you are using.

t.protein_chain = Tolower(t.protein_chain)   # protein_chain must be in a lower case

delete t.pdb_code == "3mo0"      # lig e11 2 ligands in a pocket & its pair structure is convoluted
delete t.pdb_code == "5l3e"      # lig ell 5 ligands convoluted structure
delete t.pdb_code == "5nlx"      # not relevant
delete t.pdb_code == "4or2"      # not relevant
delete t.pdb_code == "2nnh"      # lig 9cr 2 ligands in a pocket
delete t.pdb_code == "4xuh"      # lig sfi 2 ligands in a pocket
delete t.lig_name == "ret"       # pdb cis/trans all manually checked - no matching ligand pair for diff domain
delete t.lig_name == "un9"       # final retained domain were part of same doamin - manually checked

# STEP 1 - remove desolvation value below 70.
delete t.ligand_desolvation < 70.

# STEP 2 -  compare domains to ensure each "ligand" has more than one distinct InterProID
```

```
#               sort the list based on lig_name, then for each lig_name identify if it has more than one InterProID
sort(t.lig_name)
i = 1
while i <= Nof(t)
  r_lig = t.lig_name[i]
  num_lig = Nof(t.lig_name == r_lig)
  if num_lig > 1 then
    filterip = FilterIP(r_lig)
    if Type(filterip) != "integer" then
      add lig_ips filterip
    endif
    delete filterip
  endif
  i = i + num_lig
  delete r_lig
endwhile
t = lig_ips
delete lig_ips

# STEP 3 - remove duplicate PDB structures since some PDB 3D proteins structures are assigned to multiple InterProID
i = 1
while i <= Nof(t)
  r_lig = t.lig_name[i]
  t_temp = t.lig_name == r_lig
  i = i + Nof(t_temp)

  a = 1
  while a <= Nof(t_temp)
    sort(t_temp.pdb_code)
    r_pdb = t_temp.pdb_code[a]
    if Nof(t_temp.pdb_code == r_pdb) != 1 then
      delete t_temp[a]
    endif
    a = Index(t_temp.pdb_code r_pdb) + 1
    delete r_pdb
  endwhile

  if Nof(t_temp) > 1 then
    add lig_ips t_temp
  endif
  delete r_lig, t_temp
endwhile

delete t
```

<u>2- ICM conversion</u>

```
errorAction = 3

# Purpose: To generate biomolecules and save ICM objects (.ob) of PDB structures
# Input: lig_ips (table)
# Output: N/A (ICM objects saved in the folder specified by the variable "save_location")

# Prerequisite: set directory to the variable "pdb_special_cases_location" and "save_location"

#------------------------------------------------------------------------------------------------------------
# INPUT VARIABLES
t = lig_ips

# Input - directory: ensure that your directory to a specific folder ends with "\\"
pdb_special_cases_location = ""   # directory where edited PDB files are saved
save_location = ""        # directory where ICM objects will be saved
#------------------------------------------------------------------------------------------------------------

#------------------------------------------------------------------------------------------------------------
function TrimStructure( S )
# Purpose - to prepare ICM objects of PDB 3D protein structures for further analysis
# ARGUMENT:
# S[1] is pdb code
# S[2] is lig_name (with protein_chain added)
# S[3] is protein_chain

  # STEP 1 - read PDB structure
  if S[1] == "4c9w" then #read the structure where the ligand is already edited
```

```
    openFile pdb_special_cases_location + "4c9w_vgh.ob" 0 yes no no no " append"
    unds
  elseif S[1] == "5f6b" then
    openFile pdb_special_cases_location + "5f6b_ret.ob" 0 yes no no no " append"
    unds
  elseif S[1] == "5opc" then
    openFile pdb_special_cases_location + "5opc_a1z.ob" 0 yes no no no " append"
    unds
  else
    read pdb S[1]
  endif

  # STEP 2 - generate biomolecule
  makeBioMT a_$S[1]. [1] no
  if (Nof(a_*.) == 2) delete a_1.
  rename a_1. Name(Name(S[1] simple),object)

  # STEP 3 - trim structure
  delete a_$S[1].!$S[3]* #delete everything except for the molecules and ligands in the protein chain
  s = Res(Sphere(a_$S[1].$S[2] a_$S[1].$S[3] 10.))
  delete a_$S[1].$S[3]/* & !s
  convertObject Obj(a_$S[1].) 1==1 yes yes yes yes no yes ""+( 1==2 ? "water=tight ":"" )+( no ? "tautomer ":"" )
  delete S, s
endfunction
#------------------------------------------------------------------------------------------------------------------

# MAIN SCRIPT STARTS:

for i = 1, Nof(t)

  if Nof(a_*.) != 0 then
    delete a_*.
  endif
  cpdb = t.pdb_code[i]//t.protein_chain[i] + t.lig_name[i]//t.protein_chain[i]
  TrimStructure(cpdb)
  file_name = cpdb[1] + "_" + cpdb[2]
  write object auto a_$cpdb[1]. save_location+file_name + ".ob"
  delete cpdb, file_name

endfor
delete save_location, t
```

<u>3-RMSD</u>

```
errorAction = 3

# Purpose: To perform pairwise comparisons of PDB 3D protein structures co-crystalized with same ligands
#                                   by calculating RMSD values between the two structures.
# Input: lig_ips (table)
# Output: lig_rmsd (table); lig_rmsdcf (table)

# Prerequisite: set directory to the variable "icmobj_location"

#-----------------------------------------------------------------------------------------------------------
# INPUT VARIABLES
t = lig_ips

icmobj_location = ""    # use save_location specified in the script s2_bicm_conversion
cut_off = 9.5           # cut-off value for RMSD values
#-----------------------------------------------------------------------------------------------------------

# MAIN SCRIPT STARTS:

add header lig_rmsd "A new table" name = "title"
add column lig_rmsd Sarray(0) name = "lig_name"
add column lig_rmsd Rarray(0) name = "rmsd"
add column lig_rmsd Sarray(0) name = "pdb1"
add column lig_rmsd Sarray(0) name = "up_name1"
add column lig_rmsd Sarray(0) name = "ip_code1"
add column lig_rmsd Sarray(0) name = "pdb2"
add column lig_rmsd Sarray(0) name = "up_name2"
add column lig_rmsd Sarray(0) name = "ip_code2"

i = 1
```

```
while i < Nof(t) + 1
  r_lig = t.lig_name[i]
  t_temp = t.lig_name == r_lig

  for a = 1, Nof(t_temp) - 1
    if Nof(a_*.) != 0 then
      delete a_*.
    endif
    lig1 = t_temp.protein_chain[a] + r_lig
    pdb1 = t_temp.pdb_code[a]//lig1//t_temp.protein_chain[a]
    file_name1 = pdb1[1] + "_" + pdb1[2]
    openFile icmobj_location + file_name1 + ".ob" 0 yes no no no " append"
    unds

    for b = a + 1, Nof(t_temp)
      if Nof(a_*.) != 1 then
        delete a_*. & !a_$pdb1[1].
      endif
      lig2 = t_temp.protein_chain[b] + r_lig
      pdb2 = t_temp.pdb_code[b]//lig2//t_temp.protein_chain[b]
      file_name2 = pdb2[1] + "_" + pdb2[2]
      openFile icmobj_location + file_name2 + ".ob" 0 yes no no no " append"
      unds
      rmsd_val = Rmsd(a_$pdb1[1].$pdb1[3] a_$pdb2[1].$pdb2[3] align) #calculate RMSD
      add lig_rmsd
      lig_rmsd.lig_name[Nof(lig_rmsd)] = r_lig
      lig_rmsd.rmsd[Nof(lig_rmsd)] = rmsd_val
      lig_rmsd.pdb1[Nof(lig_rmsd)] = t_temp.pdb_code[a]
      lig_rmsd.up_name1[Nof(lig_rmsd)] = t_temp.up_name[a]
      lig_rmsd.ip_code1[Nof(lig_rmsd)] = t_temp.ip_code[a]
      lig_rmsd.pdb2[Nof(lig_rmsd)] = t_temp.pdb_code[b]
      lig_rmsd.up_name2[Nof(lig_rmsd)] = t_temp.up_name[b]
      lig_rmsd.ip_code2[Nof(lig_rmsd)] = t_temp.ip_code[b]
      delete sequence #due to align
      delete lig2, pdb2, rmsd_val, file_name2
    endfor #for b

    delete lig1, pdb1, file_name1
  endfor #for a

  i = i + Nof(t_temp)
  delete t_temp
endwhile

lig_rmsdcf = lig_rmsd.rmsd >= cut_off

delete t, icmobj_location, cut_off
```

<u>4- RMSD_lig_PDB</u>

```
errorAction = 3

# Purpose 1: To identify PDB 3D protein structures that meets the following criteria:
#           1) Co-crystalized with drug-like ligand (input data)
#           2) Ligand desolvation value of PDB 3D protein structures are equal or
#                     greater than 70.% (s1_data_filter)
#           2) Co-crystalized ligand of PDB 3D protein structures are co-crystalized
#                     with more than one domain (InterPro ID)(s1_data_filter)
#           4) RMSD value above cut-off (s3_rmsd)
# Purpose 2: To include protein chain, protein name, and InterPro ID for those identified
#           proteins into a single organized table

# Input: lig_rmsdcf (table); lig_ips (table)
# Output: lig_pdb (table)

#-----------------------------------------------------------------------------------------------
# INPUT VARIABLES
t = lig_rmsdcf
t_ref = lig_ips
#-----------------------------------------------------------------------------------------------

# MAIN SCRIPT STARTS:

add header lig_pdb "A new table" name = "title"
```

```
add column lig_pdb Sarray(0) name = "lig_name"
add column lig_pdb Sarray(0) name = "pdb_code"
add column lig_pdb Sarray(0) name = "protein_chain"
add column lig_pdb Sarray(0) name = "up_name"
add column lig_pdb Sarray(0) name = "ip_code"

sort(t.lig_name)
i = 1
while i < Nof(t) + 1

  ref_lig = t.lig_name[i]
  t_temp = t.lig_name == ref_lig
  for a = 1, Nof(t_temp)
    if Index(lig_pdb.pdb_code t_temp.pdb1[a]) == 0 then
      ii = Index(t_ref.pdb_code t_temp.pdb1[a])
      add lig_pdb
      lig_pdb.lig_name[Nof(lig_pdb)] = ref_lig
      lig_pdb.pdb_code[Nof(lig_pdb)] = t_temp.pdb1[a]
      lig_pdb.protein_chain[Nof(lig_pdb)] = t_ref.protein_chain[ii]
      lig_pdb.up_name[Nof(lig_pdb)] = t_ref.up_name[ii]
      lig_pdb.ip_code[Nof(lig_pdb)] = t_ref.ip_code[ii]
      delete ii
    endif
    if Index(lig_pdb.pdb_code t_temp.pdb2[a]) == 0 then
      ii = Index(t_ref.pdb_code t_temp.pdb2[a])
      add lig_pdb
      lig_pdb.lig_name[Nof(lig_pdb)] = ref_lig
      lig_pdb.pdb_code[Nof(lig_pdb)] = t_temp.pdb2[a]
      lig_pdb.protein_chain[Nof(lig_pdb)] = t_ref.protein_chain[ii]
      lig_pdb.up_name[Nof(lig_pdb)] = t_ref.up_name[ii]
      lig_pdb.ip_code[Nof(lig_pdb)] = t_ref.ip_code[ii]
      delete ii
    endif
  endfor
  i = i + Nof(t_temp)
  delete ref_lig, t_temp

endwhile

delete t, t_ref
```

<u>5- Lig-H</u>

```
errorAction = 3

# Purpose: Generate a template table for the ligand for each pdb_code, where hydrogen atoms of the ligands
#                are identified, organized, and saved into a table (tt)
# Input: lig_pdb (table)
# Output: lig_h (table)

# Prerequisite: set directory to the variable "icmobj_location" and "save_location"

#------------------------------------------------------------------------------------------------------------
# INPUT VARIABLE
t = lig_pdb

# Input - directory: ensure that your directory to a specific folder ends with "\\"
icmobj_location = ""    # use save_location specified in the script s2_bicm_conversion
save_location = ""        # directory where template table "tt" will be saved
#------------------------------------------------------------------------------------------------------------

# MAIN SCRIPT STARTS:

for i = 1, Nof(t)

  add header tt "A new table" name = "title"
  add column tt Sarray(0) name = "lig_name"
  add column tt Sarray(0) name = "pdb_code"
  add column tt Sarray(0) name = "lig_h"
  add column tt Sarray(0) name = "bound_atom"

  if Nof(a_*.) != 0 then
    delete a_*.
  endif
```

```
  cpdb = t.pdb_code[i]//t.protein_chain[i]+t.lig_name[i]//t.protein_chain[i]
  file_name = cpdb[1] + "_" + cpdb[2]
  openFile icmobj_location + file_name + ".ob" 0 yes no no no " append" #TEST
  unds

  #STEP 1 - make a table for all hydrogen atoms
  lig_h = Name(a_$cpdb[1].$cpdb[2]//h*) #identify hydrogen atoms of the ligand
  for a = 1, Nof(lig_h)
    add tt
    tt.lig_name[Nof(tt)] = t.lig_name[i]
    tt.pdb_code[Nof(tt)] = cpdb[1]
    tt.lig_h[Nof(tt)] = lig_h[a]
    bound_atom = Name(Next(a_$cpdb[1].$cpdb[2]//$lig_h[a] bond) & !a_$cpdb[1].$cpdb[2]//vt*)
    sort(bound_atom) #necessary because of "vt"
    tt.bound_atom[Nof(tt)] = bound_atom[1]
    delete bound_atom
  endfor

  #STEP 2 - identify if they are hydrogen atoms of a methyl group
  add column tt Sarray(Nof(tt)) name = "methyl"
  a = 1
  while a < Nof(tt) + 1
    ref_atom = tt.bound_atom[a]
    t_temp = tt.bound_atom == ref_atom
    if Nof(t_temp) == 3 then
      tt.methyl[a] = "yes"
      tt.methyl[a+1] = "yes"
      tt.methyl[a+2] = "yes"
    endif
    a = a + Nof(t_temp)
    delete t_temp
  endwhile

  #STEP 3 - save the final table tt to lig_energy/lig_h
  write binary tt save_location + file_name + ".icb"
  delete cpdb, file_name, lig_h, tt

endfor

delete icmobj_location, save_location, t
```

<u>6- Energy Calc.</u>

```
errorAction=3
default_terms = "vw,14,hb,el,to,ss" # default energy terms

# Purpose: To calculate the initial energies of the ligands and energies of computationally methylated ligands
#          co-crystalized with 3D protein structures "round" number of times.

# Input: lig_pdb (table)
# Output: update files in lig_h and table lig_initial

# Prerequisite: set directory to the variable "icmobj_location", "template_location" and "save_location"

#-----------------------------------------------------------------------------------------------
# INPUT VARIABLE
t = lig_pdb
round = 3 # number of rounds (n) for energy calculation - 3 rounds used in my analysis

# Input - directory: ensure that your directory to a specific folder ends with "\\"
icmobj_location = ""    # use save_location specified in the script s2_bicm_conversion
template_location = ""                  # use save_location specified in the script s5_lig_h
save_location = ""       # directory where template table "tt" will be saved
#-----------------------------------------------------------------------------------------------

#-----------------------------------------------------------------------------------------------
function DistCalculation( S )
#The function calculates and returns the distance before and after adding the methly group
#S=pdb_code//ligand name(protein chain+lig_name)//protein_chain//selected hydrogen atom
#S[4] is the selected hydrogen atom to be replaced with a methyl group

  cp a_$S[1]. "tmpObj"
  set object a_tmpObj.
```

```
  h = a_tmpObj.$S[2]//$S[4]
  min_dist1 = Min(Min(Distance( Xyz(h & !a_tmpObj.$S[2]//vt*), Xyz(a_tmpObj.$S[3]//*) )))
  modifyGroupSmiles h "C*|1.40,-0.00,0.00,0.00" no no no
  min_dist2 = Min(Min(Distance( Xyz(as_graph & !a_tmpObj.$S[2]//vt*), Xyz(a_tmpObj.$S[3]//*) )))
  delete a_tmpObj.
  delete S
  return String(min_dist1)//String(min_dist2)

endfunction

function EnergyDistCalculation( S )
#The function calculates and returns the energy vale after replacing a selected hydrogen atom with a methyl group
#S = pdb_code//ligand name(protein chain+lig_name)//protein_chain//selected hydrogen atom
#S[4] is the selected hydrogen atom to be replaced with a methyl group

  cp a_$S[1]. "tmpObj"
  set object a_tmpObj.
  h = a_tmpObj.$S[2]//$S[4]
  min_dist1 = Min(Min(Distance( Xyz(h & !a_tmpObj.$S[2]//vt*), Xyz(a_tmpObj.$S[3]//*) )))
  modifyGroupSmiles h "C*|1.40,-0.00,0.00,0.00" no no no
  set terms "tz"
  tzWeight = 200. #default is 1. - high tzWeight is required
  find molecule sstructure tether a_1.$S[2] a_tmpObj.$S[2] #template first
  minimize tether v_tmpObj.$S[2]//* 500
  mncallsMC = 10000
  montecarlo v_tmpObj.$S[2]//*
  minimize v_tmpObj.$S[2]//* 1000

  # DISTANCE AFTER MINIMIZATION
  min_dist2 = Min(Min(Distance( Xyz(as_graph & !a_tmpObj.$S[2]//vt*), Xyz(a_tmpObj.$S[3]//*) )))

  # ENERGY AFTER MINIMIZATION-eMethod2 & eMethod3
  vwMethod = 3 #old soft
  show ey a_tmpObj.$S[2] "vw,14,hb,el,to,ss"
  energy_val2 = Energy("vw,14,hb,el,to,ss")
  energy_val3 = Energy("vw,14")

  # RETURN TO DEFAULT SETTINGS
  vwMethod = 1 #default "exact"
  set terms only default_terms
  tzWeight = 1.

  delete a_tmpObj.
  delete S
  return String(energy_val2)//String(energy_val3)//String(min_dist1)//String(min_dist2)

endfunction
#---------------------------------------------------------------------------------------------------

# MAIN SCRIPT STARTS:

for n = 1, round

  if n == 1 then
    add column t Rarray(Nof(t)) name = "resolution"
  endif

  idistn = "idist_" + String(n)
  imdistn = "imdist_" + String(n)
  ieMethod2n = "ieMethod2_" + String(n)
  ieMethod3n = "ieMethod3_" + String(n)
  add column t Rarray(Nof(t)) name = idistn    # minimum distance before minimization
  add column t Rarray(Nof(t)) name = imdistn# minimum distance after minimization
  add column t Rarray(Nof(t)) name = ieMethod2n
  add column t Rarray(Nof(t)) name = ieMethod3n

  for i = 1, Nof(t)
    if Nof(a_*.) != 0 then
      delete a_*.
    endif
    cpdb = t.pdb_code[i]//t.protein_chain[i]+t.lig_name[i]//t.protein_chain[i]
    file_name = cpdb[1] + "_" + cpdb[2]
    openFile icmobj_location + file_name + ".ob" 0 yes no no no " append" #icm structure
```

```
  unds
  if n == 1 then
    openFile template_location + file_name + ".icb" #tt - template table
    t.resolution[i] = Resolution(a_$cpdb[1].)
  else
    openFile save_location + file_name + ".icb" #tt - table saved from previous round
  endif

  #STEP 1 - calculate distance for each hydrogen atom following modification (no minimization)
  distn = "dist_" + String(n)
  add column tt Rarray(Nof(tt)) name = idistn
  add column tt Rarray(Nof(tt)) name = distn
  for a = 1, Nof(tt)
    arg = cpdb//tt.lig_h[a]
    d_result = DistCalculation(arg)
    tt.$idistn[a] = Real(d_result[1])
    tt.$distn[a] = Real(d_result[2])
    delete arg, d_result
  endfor

  #STEP 2 - calculate initial energy and distance & record it to t (before and after mnimization)
  t.$idistn[i] = Min(Min(Distance( Xyz(a_$cpdb[1].$cpdb[2]//!vt*), Xyz(a_$cpdb[1].$cpdb[3]//!vt*) )))
  cp a_$cpdb[1].$cpdb[2] "lig_template"
  set object a_$cpdb[1].
  set term "tz"
  tzWeight = 1.
  find molecule sstructure tether a_lig_template.$cpdb[2] a_$cpdb[1].$cpdb[2] #template first
  minimize v_$cpdb[1].$cpdb[2]//* 1000
  delete a_lig_template.
  t.$imdistn[i] = Min(Min(Distance( Xyz(a_$cpdb[1].$cpdb[2]//!vt*), Xyz(a_$cpdb[1].$cpdb[3]//!vt*) )))
  vwMethod = 3 #old soft
  show ey a_$cpdb[1].$cpdb[2] "vw,14,hb,el,to,ss"
  i_energy2 = Energy("vw,14,hb,el,to,ss")
  i_energy3 = Energy("vw,14")

  # return to default settings - omit tzWeight (already default value)
  vwMethod = 1 #default "exact"
  set terms only default_terms

  t.$ieMethod2n[i] = Real(i_energy2)
  t.$ieMethod3n[i] = Real(i_energy3)

  # STEP 3 - calculate energy and distance after mini for each hydrogen atom following modification
  mdistn = "mdist_" + String(n)
  eMethod2n = "eMethod2_" + String(n)
  eMethod3n = "eMethod3_" + String(n)
  add column tt Rarray(Nof(tt)) name = imdistn
  add column tt Rarray(Nof(tt)) name = mdistn
  add column tt Rarray(Nof(tt)) name = eMethod2n
  add column tt Rarray(Nof(tt)) name = eMethod3n

  for a = 1, Nof(tt)
    arg = cpdb//tt.lig_h[a]
    d_result = DistCalculation(arg)
    ed_result = EnergyDistCalculation(arg)
    tt.$eMethod2n[a] = Real(ed_result[1])
    tt.$eMethod3n[a] = Real(ed_result[2])
    tt.$imdistn[a] = Real(ed_result[3])
    tt.$mdistn[a] = Real(ed_result[4])
    delete arg, ed_result
  endfor

  #STEP 4 - save tt and t(update)
  l_confirm = no
  write binary tt save_location + file_name + ".icb"
  write binary t save_location + "lig_initial.icb"
  l_confirm = yes
  delete cpdb, file_name, i_energy2, i_energy3, tt
  delete eMethod2n, eMethod3n, distn, mdistn
 endfor #for i
 delete ieMethod2n, ieMethod3n, idistn, imdistn

endfor #for n
```

lig_initial = t
delete t, round, icmobj_location, template_location, save_location

<u>7- Energy val-AS</u>

```
errorAction=3

# Purpose: To caclulate average and standard deviation of PDB 3D protein structures' energy values

# Input: lig_initial (table)
# Output: Add average and standard deviation values for each tt in pdb and table lig_initial

#-----------------------------------------------------------------------------------------------------
# INPUT VARIABLES
t = lig_initial
round = 3   # use the identical value as the variable "round" in the script s6_energycal_n

# Input - directory: ensure that your directory to a specific folder ends with "\\"
table_location = ""       # use save_location specified in the script s6_energycal_n
#-----------------------------------------------------------------------------------------------------

# MAIN SCRIPT STARTS:

add column t Rarray(Nof(t)) name = "idist_A"                    # A stands for average
add column t Rarray(Nof(t)) name = "idist_S"# S stands for standard deviation
add column t Rarray(Nof(t)) name = "imdist_A"
add column t Rarray(Nof(t)) name = "imdist_S"
add column t Rarray(Nof(t)) name = "ieMethod2_A"
add column t Rarray(Nof(t)) name = "ieMethod2_S"
add column t Rarray(Nof(t)) name = "ieMethod3_A"
add column t Rarray(Nof(t)) name = "ieMethod3_S"

for i = 1, Nof(t)

  cpdb = t.pdb_code[i]//t.protein_chain[i] + t.lig_name[i]//t.protein_chain[i]
  file_name = cpdb[1] + "_" + cpdb[2]
  openFile table_location + file_name + ".icb" #0 yes no no no "append" #tt

  add column tt Rarray(Nof(tt)) name = "idist_A"
  add column tt Rarray(Nof(tt)) name = "idist_S"
  add column tt Rarray(Nof(tt)) name = "dist_A"
  add column tt Rarray(Nof(tt)) name = "dist_S"
  add column tt Rarray(Nof(tt)) name = "imdist_A"
  add column tt Rarray(Nof(tt)) name = "imdist_S"
  add column tt Rarray(Nof(tt)) name = "mdist_A"
  add column tt Rarray(Nof(tt)) name = "mdist_S"
  add column tt Rarray(Nof(tt)) name = "eMethod2_A"
  add column tt Rarray(Nof(tt)) name = "eMethod2_S"
  add column tt Rarray(Nof(tt)) name = "eMethod3_A"
  add column tt Rarray(Nof(tt)) name = "eMethod3_S"
  add column tt Rarray(Nof(tt)) name = "deMethod2"
  add column tt Rarray(Nof(tt)) name = "deMethod3"

  # STEP 1 - calculate average and sd for initial values: ieMethod2, ieMethod3, and idist
  for n = 1, round

    idistn = "idist_" + String(n)
    imdistn = "imdist_" + String(n)
    ieMethod2n = "ieMethod2_" + String(n)
    ieMethod3n = "ieMethod3_" + String(n)
    if n == 1 then
      idistn_val = Real(t.$idistn[i])
      imdistn_val = Real(t.$imdistn[i])
      ieMethod2n_val = Real(t.$ieMethod2n[i])
      ieMethod3n_val = Real(t.$ieMethod3n[i])
    else
      idistn_ov = idistn_val
      imdistn_ov = imdistn_val
      ieMethod2n_ov = ieMethod2n_val
      ieMethod3n_ov = ieMethod3n_val
      delete idistn_val, imdistn_val, ieMethod2n_val, ieMethod3n_val
      idistn_val = idistn_ov//Real(t.$idistn[i])
      imdistn_val = imdistn_ov//Real(t.$imdistn[i])
```

```
    ieMethod2n_val = ieMethod2n_ov//Real(t.$ieMethod2n[i])
    ieMethod3n_val = ieMethod3n_ov//Real(t.$ieMethod3n[i])
    delete idistn_ov, imdistn_ov, ieMethod2n_ov, ieMethod3n_ov
  endif
  delete idistn, imdistn, ieMethod2n, ieMethod3n

endfor #for n

t.idist_A[i] = Sum(idistn_val)/Real(round)
t.idist_S[i] = Rmsd(idistn_val)
t.imdist_A[i] = Sum(imdistn_val)/Real(round)
t.imdist_S[i] = Rmsd(imdistn_val)
t.ieMethod2_A[i] = Sum(ieMethod2n_val)/Real(round)
t.ieMethod2_S[i] = Rmsd(ieMethod2n_val)
t.ieMethod3_A[i] = Sum(ieMethod3n_val)/Real(round)
t.ieMethod3_S[i] = Rmsd(ieMethod3n_val)
delete idistn_val, imdistn_val, ieMethod2n_val, ieMethod3n_val

# STEP 2 - calculate average and sd for each row in tt
for a = 1, Nof(tt)

  for n = 1, round
    idistn = "idist_" + String(n)
    distn = "dist_" + String(n)
    imdistn = "imdist_" + String(n)
    mdistn = "mdist_" + String(n)
    eMethod2n = "eMethod2_" + String(n)
    eMethod3n = "eMethod3_" + String(n)

    if n == 1 then
      idistn_val = Real(tt.$idistn[a])
      distn_val = Real(tt.$distn[a])
      imdistn_val = Real(tt.$imdistn[a])
      mdistn_val = Real(tt.$mdistn[a])
      eMethod2n_val = Real(tt.$eMethod2n[a])
      eMethod3n_val = Real(tt.$eMethod3n[a])
    else
      idistn_ov = idistn_val
      distn_ov = distn_val
      imdistn_ov = imdistn_val
      mdistn_ov = mdistn_val
      eMethod2n_ov = eMethod2n_val
      eMethod3n_ov = eMethod3n_val
      delete idistn_val, distn_val, imdistn_val, mdistn_val, eMethod2n_val, eMethod3n_val
      idistn_val = idistn_ov//Real(tt.$idistn[a])
      distn_val = distn_ov//Real(tt.$distn[a])
      imdistn_val = imdistn_ov//Real(tt.$imdistn[a])
      mdistn_val = mdistn_ov//Real(tt.$mdistn[a])
      eMethod2n_val = eMethod2n_ov//Real(tt.$eMethod2n[a])
      eMethod3n_val = eMethod3n_ov//Real(tt.$eMethod3n[a])
      delete idistn_ov, distn_ov, imdistn_ov, mdistn_ov, eMethod2n_ov, eMethod3n_ov
    endif

    delete idistn, distn, imdistn, mdistn, eMethod2n, eMethod3n
  endfor #for n

  tt.idist_A[a] = Sum(idistn_val)/Real(round)
  tt.idist_S[a] = Rmsd(idistn_val)
  tt.dist_A[a] = Sum(distn_val)/Real(round)
  tt.dist_S[a] = Rmsd(distn_val)
  tt.imdist_A[a] = Sum(imdistn_val)/Real(round)
  tt.imdist_S[a] = Rmsd(imdistn_val)
  tt.mdist_A[a] = Sum(mdistn_val)/Real(round)
  tt.mdist_S[a] = Rmsd(mdistn_val)
  tt.eMethod2_A[a] = Sum(eMethod2n_val)/Real(round)
  tt.eMethod2_S[a] = Rmsd(eMethod2n_val)
  tt.eMethod3_A[a] = Sum(eMethod3n_val)/Real(round)
  tt.eMethod3_S[a] = Rmsd(eMethod3n_val)
  delete idistn_val, distn_val, imdistn_val, mdistn_val, eMethod2n_val, eMethod3n_val

  tt.deMethod2[a]=tt.eMethod2_A[a]-t.ieMethod2_A[i]
  tt.deMethod3[a]=tt.eMethod3_A[a]-t.ieMethod3_A[i]
```

```
  endfor #for a

  # STEP 3 - save the updated tt
  l_confirm = no
  write binary tt table_location + file_name + ".icb"
  write binary t table_location + "lig_initial.icb"
  l_confirm = yes
  delete cpdb, file_name, tt

endfor #for i

lig_initial = t
delete t, table_location
```

8- Remove PDB-eM3

```
errorAction=3

# Purpose: To remove PDB 3D protein structures with initial energy values greater
#          than 0 kcal/mol calculated based on eMethod3 ("vw,14")

# Input: lig_initial (table); lig_rmsdcf (table)
# Output: dlig_initial (table); dlig_rmsdcf (table)

#-----------------------------------------------------------------------------------------------------------
# INPUT VARIABLES
t = lig_initial
tt = lig_rmsdcf
ieMethod = "ieMethod3_A"
#-----------------------------------------------------------------------------------------------------------

# MAIN SCRIPT STARTS:

t_remove = t.$ieMethod > 0

for i = 1, Nof(t_remove)
  i_remove1 = Index(tt.pdb1 t_remove.pdb_code[i])
  while i_remove1 != 0
    if (i_remove1 != 0) delete tt [i_remove1]
    i_remove1 = Index(tt.pdb1 t_remove.pdb_code[i])
  endwhile
  i_remove2 = Index(tt.pdb2 t_remove.pdb_code[i])
  while i_remove2 != 0
    if (i_remove2 != 0) delete tt [i_remove2]
    i_remove2 = Index(tt.pdb2 t_remove.pdb_code[i])
  endwhile
  delete i_remove1, i_remove2
endfor

dlig_initial = t.$ieMethod<=0
dlig_rmsdcf = tt

# test
if Index(dlig_initial.pdb_code t_remove.pdb_code[1]) != 0 then
  print "ERROR IN THE SCRIPT"
endif
if Index(dlig_rmsdcf.pdb1 t_remove.pdb_code[5]) != 0 then
  print "ERROR IN THE SCRIPT"
endif
if Index(dlig_rmsdcf.pdb2 t_remove.pdb_code[3]) != 0 then
  print "ERROR IN THE SCRIPT"
endif

delete t, tt, ieMethod, t_remove
```

9- Check-IP

```
errorAction=3

# Purpose 1: To check the domain names of each 3D protein structure pairs (inserts domain_name1 and domain_name2)
# Purpose 2: If the domain names are identical between two 3D protein structure pairs, the column remove
#            will indicate "yes" (refer to NOTE)

# Input: dlig_rmsdcf (table); ip_distribution_validation (reference table)
# Output: dlig_rmsdcf_ip (table)
```

```
# ***NOTE*** - if there are rows to remove, the changes must be updated on the table "dlig_inital" as well.
#               It wasn't necessary for the purpose of this script as there were no rows that was inidicated
#               as "yes" on the column "remove".

#-----------------------------------------------------------------------------------------------------------------
# INPUT VARIABLES
t = dlig_rmsdcf
t_ref = ip_distribution_validated
#-----------------------------------------------------------------------------------------------------------------

#-----------------------------------------------------------------------------------------------------------------
# EDIT INPUT TABLE - i.e., ip_distribution_validation table missing some domain names

# EDIT 1 - add in missing IP IDs to the reference table "t_ref"
add t_ref
t_ref.Domain_ID[Nof(t_ref)]="IPR041667"
t_ref.Domain_Name[Nof(t_ref)]="Cupin-like domain 8"

add t_ref
t_ref.Domain_ID[Nof(t_ref)]="IPR041086"
t_ref.Domain_Name[Nof(t_ref)]="YAP binding domain"

t_ref.Domain_Name[Index(t_ref.Domain_ID "IPR004045")] = "Glutathione S-transferase"
t_ref.Domain_Name[Index(t_ref.Domain_ID "IPR000719")] = "Protein kinase domain"

# EDIT 2 - replace redundant IP IDs on table "t"
t.ip_code1=Replace(t.ip_code1 "IPR004046" "IPR004045") #GST
t.ip_code2=Replace(t.ip_code2 "IPR004046" "IPR004045")

t.ip_code1=Replace(t.ip_code1 "IPR001245" "IPR000719") #Protein kinase domain
t.ip_code2=Replace(t.ip_code2 "IPR001245" "IPR000719")
#-----------------------------------------------------------------------------------------------------------------

# MAIN SCRIPT STARTS:

add column t Sarray(Nof(t)) name="domain_name1"
add column t Sarray(Nof(t)) name="domain_name2"
add column t Sarray(Nof(t)) name="remove"

for i=1,Nof(t)
  if t.ip_code1[i]==t.ip_code2[i] then
    t.remove[i]="yes"
    continue
  endif
  t.domain_name1[i] = t_ref.Domain_Name[Index(t_ref.Domain_ID t.ip_code1[i])]
  t.domain_name2[i] = t_ref.Domain_Name[Index(t_ref.Domain_ID t.ip_code2[i])]
endfor

if Nof(t.remove=="yes")==0 then #NOTE: if not, then write a code to remove it from "dlig_inital" as well
  delete t.remove
endif

dlig_rmsdcf_ip = t
delete t_ref, t
```

<u>10- H-Match</u>

```
errorAction=3

# Purpose 1: To check the domain names of each 3D protein structure pairs (inserts domain_name1 and domain_name2)
# Purpose 2: If the domain names are identical between two 3D protein structure pairs, the column remove
#               will indicate "yes" (refer to NOTE)

# Input: dlig_rmsdcf (table); ip_distribution_validation (reference table)
# Output: dlig_rmsdcf_ip (table)

# ***NOTE*** - if there are rows to remove, the changes must be updated on the table "dlig_inital" as well.
#               It wasn't necessary for the purpose of this script as there were no rows that was inidicated
#               as "yes" on the column "remove".

#-----------------------------------------------------------------------------------------------------------------
# INPUT VARIABLES
t = dlig_rmsdcf
```

```
t_ref = ip_distribution_validated
#--------------------------------------------------------------------------------------------------------------------

#--------------------------------------------------------------------------------------------------------------------
# EDIT INPUT TABLE - i.e., ip_distribution_validation table missing some domain names

# EDIT 1 - add in missing IP IDs to the reference table "t_ref"
add t_ref
t_ref.Domain_ID[Nof(t_ref)]="IPR041667"
t_ref.Domain_Name[Nof(t_ref)]="Cupin-like domain 8"

add t_ref
t_ref.Domain_ID[Nof(t_ref)]="IPR041086"
t_ref.Domain_Name[Nof(t_ref)]="YAP binding domain"

t_ref.Domain_Name[Index(t_ref.Domain_ID "IPR004045")] = "Glutathione S-transferase"
t_ref.Domain_Name[Index(t_ref.Domain_ID "IPR000719")] = "Protein kinase domain"

# EDIT 2 - replace redundant IP IDs on table "t"
t.ip_code1=Replace(t.ip_code1 "IPR004046" "IPR004045") #GST
t.ip_code2=Replace(t.ip_code2 "IPR004046" "IPR004045")

t.ip_code1=Replace(t.ip_code1 "IPR001245" "IPR000719") #Protein kinase domain
t.ip_code2=Replace(t.ip_code2 "IPR001245" "IPR000719")
#--------------------------------------------------------------------------------------------------------------------

# MAIN SCRIPT STARTS:

add column t Sarray(Nof(t)) name="domain_name1"
add column t Sarray(Nof(t)) name="domain_name2"
add column t Sarray(Nof(t)) name="remove"

for i=1,Nof(t)
  if t.ip_code1[i]==t.ip_code2[i] then
    t.remove[i]="yes"
    continue
  endif
  t.domain_name1[i] = t_ref.Domain_Name[Index(t_ref.Domain_ID t.ip_code1[i])]
  t.domain_name2[i] = t_ref.Domain_Name[Index(t_ref.Domain_ID t.ip_code2[i])]
endfor

if Nof(t.remove=="yes")==0 then #NOTE: if not, then write a code to remove it from "dlig_inital" as well
  delete t.remove
endif

dlig_rmsdcf_ip = t
delete t_ref, t
```

<u>11- Lig-Energy</u>

```
errorAction = 3
default_terms = "vw,14,hb,el,to,ss"

# Purpose: To match hydorgen atom # (and associated energy values) of ligands between two PDB 3D structures of each pair

# Input: dlig_rmsdcf_ip (table); dlig_initial (reference table)
# Output: lig_energy (table)

# Prerequisite: set directory to the variable "table_location", "h_matched_tt_location" and "save_location"

#-----------------------------------------------------------------------------------------------
# INPUT VARIABLES
t = dlig_rmsdcf_ip
t_ref = dlig_initial

distn = "dist_A"           # depending on the energy column in interest (average vs. single val)
mdistn = "mdist_A"
eMethod2n = "eMethod2_A"
eMethod3n = "eMethod3_A"
deMethod2n = "deMethod2"
deMethod3n = "deMethod3"

# Input - directory: ensure that your directory to a specific folder ends with "\\"
table_location = ""      # use table_location specified in the script s7_energyval_AS
```

```
h_matched_tt_location = ""          # use folder location where tables in save_location specified in the script s10_h_match
#                                                                    are manually checked and corrected (refer to NOTE below)
save_location = ""       # directory where output table "lig_energy" will be saved

# NOTE - "h_matched_tt_location" (hydrogen atoms matched template table location) is the location in
#               which template tables of list of hydrogen atoms of the same co-crystalized ligands matched
#               between two PDB structures.
#               IF the hydrogen atoms are matched using built-in function, they are not perfect. Hence, it
#               is required to be manually checked as also written in the script s10_h_match.
#------------------------------------------------------------------------------------------------

# MAIN SCRIPT STARTS:

add column t Sarray(Nof(t)) name = "error_s7"

if Nof(a_*.) != 0 then
  delete a_*.
endif

for i = 1, Nof(t)

  index1=Index(t_ref.pdb_code t.pdb1[i])
  index2=Index(t_ref.pdb_code t.pdb2[i])
  pdb1=t.pdb1[i]//t_ref.protein_chain[index1]+t.lig_name[i]//t.lig_name[i]
  pdb2=t.pdb2[i]//t_ref.protein_chain[index2]+t.lig_name[i]//t.lig_name[i]
  delete index1,index2

  file_name1 = pdb1[1] + "_" + pdb1[2]
  file_name2 = pdb2[1] + "_" + pdb2[2]
  file_name = pdb1[2] + "_" + pdb1[1] + "_" + pdb2[1]

  openFile table_location + file_name1 + ".icb" #tt
  rename tt Name(Name("pdb1t" simple),unique)
  openFile table_location + file_name2 + ".icb" #tt
  rename tt Name(Name("pdb2t" simple),unique)
  if Nof(pdb1t) != Nof(pdb2t) then #error check
    delete index1, index2, pdb1, pdb2, pdb1t, pdb2t
    t.error_s7[i] = "yes"
    continue
  endif

  openFile h_matched_tt_location + file_name + ".icb" #tt - matched h table

  #STEP 2 - organize information based on mathcing hydrogens (tt) into lig_energy
  add header lig_energy "A new table" name = "title"
  add column lig_energy Sarray(0) name = "pdb1_h"
  add column lig_energy Sarray(0) name = "pdb2_h"
  add column lig_energy Sarray(0) name = "methyl_h"
  add column lig_energy Rarray(0) name = "pdb1_dist"
  add column lig_energy Rarray(0) name = "pdb2_dist"
  add column lig_energy Rarray(0) name = "pdb1_mdist"
  add column lig_energy Rarray(0) name = "pdb2_mdist"
  add column lig_energy Rarray(0) name = "pdb1_eMethod2"
  add column lig_energy Rarray(0) name = "pdb2_eMethod2"
  add column lig_energy Rarray(0) name = "pdb1_eMethod3"
  add column lig_energy Rarray(0) name = "pdb2_eMethod3"

  a = 1
  sort(pdb1t.bound_atom)
  while a < Nof(pdb1t) + 1 #while h is less than number of hydrogen atoms

    add lig_energy
    if pdb1t.methyl[a] == "yes" then #if methyl then find and use the lowest energy val
      lig_energy.methyl_h[Nof(lig_energy)] = "yes"
      t_temp1 = pdb1t.bound_atom == pdb1t.bound_atom[a]
      t_temp2 = pdb2t.bound_atom == tt.pdb2_atom[Index(tt.pdb1_atom pdb1t.bound_atom[a])]

      pdb1_h = t_temp1.lig_h[1] + "/" + t_temp1.lig_h[2] + "/" + t_temp1.lig_h[3]
      h1_index = Index(t_temp2.lig_h tt.pdb2_atom[Index(tt.pdb1_atom t_temp1.lig_h[1])])
      h2_index = Index(t_temp2.lig_h tt.pdb2_atom[Index(tt.pdb1_atom t_temp1.lig_h[2])])
      h3_index = Index(t_temp2.lig_h tt.pdb2_atom[Index(tt.pdb1_atom t_temp1.lig_h[3])])
      pdb2_h = t_temp2.lig_h[h1_index] + "/" + t_temp2.lig_h[h2_index] + "/" + t_temp2.lig_h[h3_index]
```

```
      highest_dist = Sort(t_temp1.$distn)[Nof(t_temp1)]//Sort(t_temp2.$distn)[Nof(t_temp2)]
      highest_mdist = Sort(t_temp1.$mdistn)[Nof(t_temp1)]//Sort(t_temp2.$mdistn)[Nof(t_temp2)]
      lowest_eMethod2 = Sort(t_temp1.$eMethod2n)[1]//Sort(t_temp2.$eMethod2n)[1]
      lowest_eMethod3 = Sort(t_temp1.$eMethod3n)[1]//Sort(t_temp2.$eMethod3n)[1]

      pdb1_info = highest_dist[1]//highest_mdist[1]//lowest_eMethod2[1]//lowest_eMethod3[1]
      pdb2_info = highest_dist[2]//highest_mdist[2]//lowest_eMethod2[2]//lowest_eMethod3[2]
      a = a + 3
      delete t_temp1, t_temp2, h1_index, h2_index, h3_index
      delete highest_dist, highest_mdist, lowest_eMethod2, lowest_eMethod3
    else
      pdb1_h = pdb1t.lig_h[a]
      pdb1_info = pdb1t.$distn[a]//pdb1t.$mdistn[a]//pdb1t.$eMethod2n[a]//pdb1t.$eMethod3n[a]
      pdb2_h = tt.pdb2_atom[Index(tt.pdb1_atom pdb1_h)]
      pdb2_index = Index(pdb2t.lig_h pdb2_h)
      pdb2_info = pdb2t.$distn[pdb2_index]//pdb2t.$mdistn[pdb2_index]//pdb2t.$eMethod2n[pdb2_index]//pdb2t.$eMethod3n[pdb2_index]
      a = a + 1
      delete pdb2_index
    endif

    lig_energy.pdb1_h[Nof(lig_energy)] = pdb1_h
    lig_energy.pdb2_h[Nof(lig_energy)] = pdb2_h
    lig_energy.pdb1_dist[Nof(lig_energy)] = pdb1_info[1]
    lig_energy.pdb2_dist[Nof(lig_energy)] = pdb2_info[1]
    lig_energy.pdb1_mdist[Nof(lig_energy)] = pdb1_info[2]
    lig_energy.pdb2_mdist[Nof(lig_energy)] = pdb2_info[2]
    lig_energy.pdb1_eMethod2[Nof(lig_energy)] = pdb1_info[3]
    lig_energy.pdb2_eMethod2[Nof(lig_energy)] = pdb2_info[3]
    lig_energy.pdb1_eMethod3[Nof(lig_energy)] = pdb1_info[4]
    lig_energy.pdb2_eMethod3[Nof(lig_energy)] = pdb2_info[4]
    delete pdb1_h, pdb2_h, pdb1_info, pdb2_info

  endwhile

  #STEP 3 - save lig_energy
  l_confirm = no
  write binary lig_energy save_location + file_name + ".icb"
  l_confirm = yes
  delete pdb1, pdb2, pdb1t, pdb2t, h_match, file_name1, file_name2, file_name, tt, lig_energy

endfor

if Nof(t.error_s7 == "yes") != 0 then
  error_s7 = t
endif

delete t, t_ref, distn, mdistn, eMethod2n, eMethod3n, save_location
```

12- Percent risk

```
errorAction = 3

# Purpose: To calculate percent risk, pcR (previously termed shared percentage (sp)).
#            "lig_energy" tables generated and saved in the script s12_lig_energy are updated.

# Input: dlig_rmsdcf_ip (table); dlig_initial (reference table)
# Output: lig_sp

# Prerequisite: set directory to the variable "table_location" and "save_location"

#-----------------------------------------------------------------------------------------
# INPUT VARIABLES
t = dlig_rmsdcf_ip
t_ref = dlig_initial

# Input - directory: ensure that your directory to a specific folder ends with "\\"
table_location = ""      # use save_location specified in the script s11_lig_energy
save_location = ""       # directory where updated "lig_energy" tables will be saved

dist_threshold=1.6      # threshold/cut-off values
eMethod2_threshold=0.
eMethod3_threshold=0.
#-----------------------------------------------------------------------------------------
```

```
# MAIN SCRIPT STARTS----------------------------------------------------------------------
add column t Rarray(Nof(t)) name = "pdb1_d_sp"           #sp (shared percentage) == pcR (percent risk)
add column t Rarray(Nof(t)) name = "pdb2_d_sp"
add column t Rarray(Nof(t)) name = "pdb1_md_sp"
add column t Rarray(Nof(t)) name = "pdb2_md_sp"
add column t Rarray(Nof(t)) name = "pdb1_eM2_sp"
add column t Rarray(Nof(t)) name = "pdb2_eM2_sp"
add column t Rarray(Nof(t)) name = "pdb1_eM3_sp"
add column t Rarray(Nof(t)) name = "pdb2_eM3_sp"
#<optional>
add column t Rarray(Nof(t)) name = "pdb1_eM2_bp"     # bp (buried percentage) == cp (constraint percentage)
add column t Rarray(Nof(t)) name = "pdb2_eM2_bp"
add column t Rarray(Nof(t)) name = "pdb1_eM3_bp"
add column t Rarray(Nof(t)) name = "pdb2_eM3_bp"

for i = 1, Nof(t)

 index1 = Index(t_ref.pdb_code t.pdb1[i])
 index2 = Index(t_ref.pdb_code t.pdb2[i])
 pdb1 = t.pdb1[i]//t_ref.protein_chain[index1] + t.lig_name[i]//t.lig_name[i]
 pdb2 = t.pdb2[i]//t_ref.protein_chain[index2] + t.lig_name[i]//t.lig_name[i]
 delete index1, index2

 file_name = pdb1[2] + "_" + pdb1[1] + "_" + pdb2[1]
 openFile table_location + file_name + ".icb"

 add column lig_energy Sarray(Nof(lig_energy)) name = "d_shared"
 add column lig_energy Sarray(Nof(lig_energy)) name = "md_shared"
 add column lig_energy Sarray(Nof(lig_energy)) name = "eM2_shared"
 add column lig_energy Sarray(Nof(lig_energy)) name = "eM3_shared"

 for a = 1, Nof(lig_energy)
  if lig_energy.pdb1_dist[a] < dist_threshold & lig_energy.pdb2_dist[a] < dist_threshold then
   lig_energy.d_shared[a] = "yes"
  endif
  if lig_energy.pdb1_mdist[a] < dist_threshold & lig_energy.pdb2_mdist[a] < dist_threshold then
   lig_energy.md_shared[a] = "yes"
  endif
  if lig_energy.pdb1_eMethod2[a] > eMethod2_threshold & lig_energy.pdb2_eMethod2[a] > eMethod2_threshold then
   lig_energy.eM2_shared[a] = "yes"
  endif
  if lig_energy.pdb1_eMethod3[a] > eMethod3_threshold & lig_energy.pdb2_eMethod3[a] > eMethod3_threshold then
   lig_energy.eM3_shared[a] = "yes"
  endif
 endfor #for a

 nof_d_shared = Real(Nof(lig_energy.d_shared == "yes"))
 nof_md_shared = Real(Nof(lig_energy.md_shared == "yes"))
 nof_eM2_shared = Real(Nof(lig_energy.eM2_shared == "yes"))
 nof_eM3_shared = Real(Nof(lig_energy.eM3_shared == "yes"))

 if Nof(lig_energy.pdb1_dist < dist_threshold) != 0 then
  t.pdb1_d_sp[i] = nof_d_shared/Real(Nof(lig_energy.pdb1_dist < dist_threshold))*100
 else
  t.pdb1_d_sp[i] = -1.
 endif
 if Nof(lig_energy.pdb2_dist < dist_threshold) != 0 then
  t.pdb2_d_sp[i] = nof_d_shared/Real(Nof(lig_energy.pdb2_dist < dist_threshold))*100
 else
  t.pdb2_d_sp[i] = -1.
 endif

 if Nof(lig_energy.pdb1_mdist < dist_threshold) != 0 then
  t.pdb1_md_sp[i] = nof_md_shared/Real(Nof(lig_energy.pdb1_mdist < dist_threshold))*100
 else
  t.pdb1_md_sp[i] = -1.
 endif
 if Nof(lig_energy.pdb2_mdist < dist_threshold) != 0 then
  t.pdb2_md_sp[i] = nof_md_shared/Real(Nof(lig_energy.pdb2_mdist < dist_threshold))*100
 else
  t.pdb2_md_sp[i] = -1.
 endif
 if Nof(lig_energy.pdb1_eMethod2 > eMethod2_threshold) != 0 then
```

```
    t.pdb1_eM2_sp[i] = nof_eM2_shared/Real(Nof(lig_energy.pdb1_eMethod2 > eMethod2_threshold))*100
    t.pdb1_eM2_bp[i] = Real(Nof(lig_energy.pdb1_eMethod2 > eMethod2_threshold))/Real(Nof(lig_energy))*100
  else
    t.pdb1_eM2_sp[i] = -1.
    t.pdb1_eM2_bp[i] = 0.
  endif
  if Nof(lig_energy.pdb2_eMethod2 > eMethod2_threshold) != 0 then
    t.pdb2_eM2_sp[i] = nof_eM2_shared/Real(Nof(lig_energy.pdb2_eMethod2 > eMethod2_threshold))*100
    t.pdb2_eM2_bp[i] = Real(Nof(lig_energy.pdb2_eMethod2>eMethod2_threshold))/Real(Nof(lig_energy))*100
  else
    t.pdb2_eM2_sp[i] = -1.
    t.pdb2_eM2_bp[i] = 0.
  endif
  if Nof(lig_energy.pdb1_eMethod3 > eMethod3_threshold) != 0 then
    t.pdb1_eM3_sp[i] = nof_eM3_shared/Real(Nof(lig_energy.pdb1_eMethod3 > eMethod3_threshold))*100
    t.pdb1_eM3_bp[i] = Real(Nof(lig_energy.pdb1_eMethod3 > eMethod3_threshold))/Real(Nof(lig_energy))*100
  else
    t.pdb1_eM3_sp[i] = -1.
    t.pdb1_eM3_sp[i] = 0.
  endif
  if Nof(lig_energy.pdb2_eMethod3 > eMethod3_threshold) != 0 then
    t.pdb2_eM3_sp[i] = nof_eM3_shared/Real(Nof(lig_energy.pdb2_eMethod3 > eMethod3_threshold))*100
    t.pdb2_eM3_bp[i] = Real(Nof(lig_energy.pdb2_eMethod3 > eMethod3_threshold))/Real(Nof(lig_energy))*100
  else
    t.pdb2_eM3_sp[i] = -1.
    t.pdb2_eM3_bp[i] = 0.
  endif

  delete nof_d_shared, nof_md_shared, nof_eM2_shared, nof_eM3_shared
  l_confirm = no
  write binary lig_energy save_location + file_name + ".icb"
  write binary t save_location + "lig_sp.icb"
  l_confirm = yes
  delete pdb1, pdb2, file_name, lig_energy

endfor

lig_sp = t
delete t, t_ref, dist_threshold, eMethod2_threshold, eMethod3_threshold
delete save_location, table_location
```
